# Appetite for change: How nutritional state and ventral striatal dopamine shape psilocybin effects on cognitive flexibility

**DOI:** 10.64898/2026.05.14.725265

**Authors:** Kyna Conn, Ngok Wong, Serenay Sunnetci, Abdallah Al Siyabi, Kaspar McCoy, Kaixin Huang, Erika Greaves, Laura K Milton, Levin Kuhlmann, Sarah Maguire, Claire J Foldi

## Abstract

Psilocybin has emerged as a promising therapeutic agent for psychiatric disorders characterised by cognitive rigidity and disrupted reward processing, including anorexia nervosa. While its pro-cognitive effects have been mechanistically probed almost exclusively through serotonin receptor subtype antagonism, the downstream contributions of dopaminergic systems to these outcomes remain poorly understood. Here, we examined how psilocybin (1.5 mg/kg) alters cognitive flexibility across multiple operant paradigms in female rats and whether ventral striatal dopamine, nutritional state or prior activity-based anorexia (ABA) exposure moderate these effects. In food-restricted rats, psilocybin improved reversal learning with an apparent temporal delay relative to our previously published *ad libitum* fed cohort. *In vivo* fiber photometry revealed that psilocybin broadly amplified nucleus accumbens core (NAc) dopamine transients time-locked to both expected and unexpected outcomes during probabilistic reversal learning across 7 days. Computational modelling identified psilocybin-specific increases in learning rate and reductions in prior value weighting, consistent with strengthened feedback-driven updating. In touchscreen paradigms, psilocybin enhanced discrimination accuracy and accelerated reversal learning acquisition when administered prior to initial discrimination, but impaired serial reversal accuracy when administered after initial discrimination. ABA exposure constrained psilocybin’s pro-cognitive effects, abolishing discrimination accuracy benefits, suggesting that metabolic disruption and stress alter the neural substrates required for psilocybin-enhanced neuroplasticity and reward learning. These findings provide the first direct evidence that psilocybin modulates striatal dopamine signalling in a behaving animal and demonstrate that current nutritional state and prior experience with pathological weight loss and reward learning moderate its cognitive effects.

## Introduction

The classical psychedelic psilocybin has emerged as a promising therapeutic agent for treatment-resistant psychiatric disorders, including depression [1], addiction [2], and anorexia nervosa (AN) [3]. Beyond its well-documented acute psychoactive effects, psilocybin demonstrates potential to enhance cognitive processes that are frequently impaired in humans, with the strongest evidence supporting enhancement of cognitive flexibility [4–6], an executive function that coordinates the ability to adaptively shift behavioural strategies in response to changing environmental demands. In clinical populations characterized by cognitive rigidity, psilocybin demonstrates the capacity to ameliorate inflexible thinking patterns in people with depression [4], and may restructure rigid maladaptive thoughts about food and feeding in people with AN [3]. Preclinical studies have shown improvements in flexible learning using set-shifting tasks [7] and reversal learning paradigms [8]. However, the effects on preclinical outcomes are not uniformly positive, with some studies reporting context- and dose-dependent impairments [9–11], while others suggest a nuanced role in which psilocybin can transiently impair learning but ultimately enhance cognitive flexibility [12]. This suggests that the effects of psilocybin on cognitive function may depend on factors such as dosing, timing, environmental conditions, and baseline cognitive or neurochemical state.

The majority of mechanistic research into the actions of psilocybin that drive improved cognitive flexibility has focused on serotonin (5-HT) signalling, in particular 5-HT2A receptor activation in cortical regions [7, 8, 13]. Indeed, we have previously shown that psilocybin acutely alters the transcription of serotonin receptors in the rat medial prefrontal cortex, an effect that was exacerbated in animals that had undergone exposure to a preclinical model of AN known as activity-based anorexia (ABA) [8]. While recent calls to examine targets of psilocybin beyond the 5-HT2A receptor are encouraging [14], the downstream contributions of dopaminergic systems to its cognitive effects remain underexplored, which is surprising given that substantial cross-talk occurs between serotonergic and dopaminergic systems [15] and the dense expression of 5-HT2A receptors on cortical pyramidal neurons that project directly to striatal regions, where stimulation triggers glutamate release that subsequently modulates striatal dopamine activity [16]. Of note, the ABA phenotype is driven by alterations in neurochemical and metabolic function, where both serotonin and dopamine system influences have been observed [17–19], including at the level of receptor binding [20]. Thus, it is plausible that baseline neurochemical and/or metabolic state can impact the way in which psilocybin acts in the brain to modify flexible learning.

Disruptions within fronto-striatal circuitry mediate the cognitive inflexibility observed in obsessive-compulsive disorder, substance use disorders, and depression [21–23]. This dysfunction is particularly pronounced in AN, where individuals demonstrate marked rigidity around food-related decisions, an inability to update maladaptive eating behaviours despite negative health consequences, and profound deficits in reward processing that perpetuate restrictive eating patterns [24]. Within this cortico-striatal architecture, the ventral striatum (specifically the nucleus accumbens; NAc) serves as a critical node integrating excitatory inputs from the prefrontal cortex (PFC), anterior cingulate cortex, and insular cortex; conveying higher-order information about goals, action outcomes, effort, and internal states; with dopaminergic projections from the ventral tegmental area to guide flexible decision-making [25]. Central to this circuitry is phasic dopaminergic signalling, which encodes reward prediction errors and provides critical teaching signals that update behavioural strategies when environmental contingencies change [26]. Human positron emitted tomography (PET) studies using raclopride displacement [27] and microdialysis experiments in anaesthetised rats [28] confirm that psilocybin increases extracellular striatal dopamine availability in the NAc. By modulating ventral striatal dopamine dynamics and outcome-related prediction errors, psilocybin my enhance feedback-driven updating to promote flexible reward learning.

Understanding whether psilocybin alters ventral striatal dopaminergic mechanisms is particularly relevant for its therapeutic application in AN, a disorder characterized by both profound cognitive rigidity and altered reward processing [29] and reinforcement learning [30]. Specific neuropsychiatric symptoms present in people with AN, such as anhedonia [31] or restrictive feeding [32], are often linked to NAc hypodopaminergia, reducing reward sensitivity and impairing adaptive behaviour. Further complicating interpretation of potential dopaminergic actions of psilocybin is the fact that energy deficits, acute food restriction or altered nutritional or metabolic state will likely be present in individuals with AN accessing psilocybin assisted psychotherapy, states that are also well known to alter dopaminergic tone and receptor availability [33–35], changes that may explain food avoidance behaviours in AN [36, 37]. However, how an individual’s current nutritional state or prior experience with pathological weight loss impacts responsiveness to psilocybin remains entirely unknown. Such an understanding may be important for understanding the heterogeneity in response profiles observed in clinical trials evaluating the efficacy of psilocybin for AN [3, 38]. Therefore, the objective of the present study was to use the ABA rat model and reversal learning paradigms to examine 1) how current energy state affects psilocybin-enhanced flexible learning, 2) whether ventral striatal dopamine release tracks with psilocybin-induced learning improvements and 3) how timing of psilocybin administration and prior experience with pathological weight loss modulate effects on cognitive flexibility. Our goal is to develop a mechanistic model of psilocybin effects relevant to its clinical application in AN and other eating disorders.

## Methods

### Animals and Housing

Young female Sprague Dawley rats (*N*=139; Monash Animal Research Platform, Clayton, VIC, Australia) were used based on their susceptibility to activity-based anorexia (ABA) and translational relevance to anorexia nervosa. Rats were group-housed and acclimated to a 12 h light/dark cycle (lights off at 1100 h) for 7 days in temperature- (22–24°C) and humidity-controlled (30–50%) conditions with *ad libitum* access to water and standard chow (Barastoc, Australia) unless undergoing food restriction. To facilitate oestrous cycle synchronisation (Whitten Effect; [39]), a pair of male rats was housed in each experimental room ≥7 days prior to experimentation. ABA modelling began at 6–7 weeks of age as previously described [8, 40, 41]; all feeding and operant experiments began at 10–12 weeks of age. For most behavioural experiments, animals were individually housed (26 W × 21 H × 47.5 D cm). Touchscreen-based (PhenoSys) experiments used group housing (5–8 per cage, 34 W x 26 H × 55 D cm) to enable RFID-sorted access to automated testing chambers [40]. Food-restricted animals were titrated to 90–95% of baseline body weight. To adhere to the 3Rs principles of animal ethics (Reduction) and minimize unnecessary animal use, *ad libitum* control data for FED3 reversal learning were drawn from our previously published dataset [8], collected in the identical facility, on the same hardware platforms, using identical light cycles, strain, sex, and testing protocols. All procedures were approved by the Monash Animal Resource Platform Ethics Committee (ERM 36800) in accordance with the Australian Code for the care and use of animals for scientific purposes. See **Supplementary Table 1** for animal numbers in each experiment.

### Activity-Based Anorexia (ABA) Model

The ABA paradigm consists of unlimited access to a running wheel and time-restricted food access. At 7 weeks, rats (*N*=17) were individually housed with a running wheel (Lafayette Instruments, IN, USA) and a removable food basket. After 7 days of wheel habituation to establish baseline running wheel activity (RWA), food access was restricted to 90 min/day at dark phase onset (1100–1230 h) for up to 10 days or until rats reached <80% baseline body weight, at which point they were refed and wheels locked. RWA was recorded using Scurry Activity Wheel Software (Lafayette Instruments, IN, USA). Rats were allowed to recovery body weight (>100% baseline) before being tested on touchscreen-based reversal learning tasks (see below). Control animals (ABA-naïve) were age-matched female rats maintained under standard *ad libitum* feeding and social housing conditions without access to running wheels, but subjected to identical handling and operant testing schedules.

### Drug Administration

Psilocybin (USONA Institute, Lot# AMS0167) was dissolved in saline and administered intraperitoneally at 1.5 mg/kg (1.0 ml/kg, 26-gauge needle), selected based on established occupancy and prior behavioural efficacy in rats; vehicle controls received 0.9% NaCl. A single dose of psilocybin was administered 18-20h prior to the first behavioural test session of the target phase across experiments; either the first session using 80:20 probabilistic contingencies, Pairwise Discrimination (PD) or reversal of reward contingencies (R1), or 90 min prior to perfusion for cFos analyses.

### Psilocybin-induced Activation of NAc Neurons

To assess whether food restriction altered psilocybin-induced activation of the NAc compared to *ad libitum* feeding, rats were euthanised with sodium pentobarbitone (Lethabarb, 150 mg/kg; Virbac, AU) and transcardially perfused with 0.9% saline (200 mL) followed by 4% paraformaldehyde in phosphate buffer (200 mL). Brains were post-fixed overnight at 4°C then transferred to 30% sucrose in phosphate buffer for 24 h. Coronal sections (35 μm) encompassing the NAc (bregma AP: +1.6 to +4.8 mm) were collected at a 1:4 series using a cryostat (CM1860; Leica Biosystems). Immunohistochemistry utilised primary (guinea pig anti-cFos; 1:3000, Synaptic Systems 226308) and secondary (goat anti-guinea pig Alexa fluor 488; 1:500, Abcam ab150185) antibodies; DAPI (1:200, Sigma-Aldrich, CAS 28718-90-3) was used for nuclear staining. Three sections per animal were imaged on a widefield microscope (Thunder Imager Live Cell & 3D Assay, Leica Microsystems, Germany) with a 10× objective, deconvolved using LIGHTNING (Leica), and analysed using a custom ImageJ macro (v1.53t; [42]) for automated quantification of DAPI+ and cFos+ puncta.

### Between-Sessions Reversal Learning

Animals were trained on open-source Feeding Experimentation Devices Version 3 (FED3 [43]) operant devices in daily sessions during the mid-dark phase exactly as previously described [8], although with shorter session times (1 h instead of 3 h) and under food restriction to maintain body weight at ∼90% of baseline. Training progressed through FR1, FR3, and FR5 across 3 days each, until >80% accuracy at the target port was achieved. Rats received sucrose reward pellets (20mg, AS5TUT; Test Diet, CA, USA) to positively reinforce correct actions. Psilocybin or saline was administered following the final FR5 session, and the target port was reversed the following day. Reversal learning was assessed across three daily 1 h sessions, with groups counterbalanced for side bias.

### Stereotaxic Surgery and Viral Preparation

Rats were anaesthetised with isoflurane (5% induction, 2.5% maintenance; 0.5 L/min flow rate) and positioned in a stereotaxic frame. A glass micropipette (Drummond Scientific; #5-000-1001-X10) was used to infuse 400 nL of AAV dopamine biosensor (pAAV-hsyn-GRAB_DA2m; Addgene #140553) unilaterally into the NAc core (AP: +1.70; ML: ±1.50; DV: −6.2 mm from bregma) at 80 nL/min, with a 5-min post-infusion wait. A fibre optic cannula (200 μm core, 7.5 mm length, NA 0.37, 1.25 mm ferrule; RWD Systems) was implanted at the same coordinates and secured with stainless steel skull screws and dental cement. Rats recovered for ≥7 days prior to habituation. See **Supplementary Methods** for full surgical details.

### Within-session Probabilistic Reversal Learning (FED) and In Vivo Fibre Photometry

Animals underwent daily 1 h FED3 sessions under food restriction. Following 1 week of bilateral nose-poke training, rats were assessed on a deterministic 100:0 reversal paradigm (reversal triggered after 10 consecutive correct responses) for 2 weeks, concurrent with fibre photometry habituation (Neurophotometrics FP3002). Drug was administered following the final deterministic session. Probabilistic reversal learning (80:20 contingencies) was then conducted for 7 post-administration days paired with dopamine recordings in the NAc. Full methodological details available in **Supplementary Methods**.

Fluorescence signals were time-locked to FED3 nose-poke and reward events, and acquired via Bonsai (Version 2.7.1; [44]). Dual LEDs (470 nm and 415 nm) were alternately pulsed at 40 Hz; 470 nm reflected GRAB-DA sensor excitation and 415 nm served as an isosbestic control. Raw fluorescence was first baseline-corrected via ΔF/F using the fitted isosbestic control and then normalized by converting to z-scores relative to a pre-event baseline period. This expresses dopamine transients as standardized deviations from baseline activity, enabling comparison across trials and subjects. Behavioural analyses included *n*=8–9 animals per group; animals failing to achieve ≥3 reversals/session during deterministic training were excluded *a priori*. Photometry analyses used n=4 per group following exclusion for fibre misplacement or excessive noise, verified by post hoc immunohistochemistry confirming GRAB-DA expression and fibre placement in the NAc.

### Computational Modelling

To characterise latent decision processes during reversal learning, we fitted a Rescorla–Wagner reinforcement-learning (RL) model and an active inference (AI) model to the trial-by-trial behavioural data. Both models were fit separately to each rat-session using the same extracted choice and outcome arrays. Model comparison was implemented using the Bayesian Information Criterion (BIC), calculated separately for each fitted rat-session as BIC=kln(n)+2 NLL, where k is the number of free parameters, n is the number of trials, and NLL is the negative log-likelihood of the fitted model [45]. Lower BIC values indicate a better trade-off between goodness of fit and model complexity. Session-wise BIC values were then compared between the RW and AI models and additionally summarised within animal for rat-level comparison. See **Supplementary Methods** for extended details of model parameters.

### Pairwise Discrimination and Reversal Learning: Automated Touchscreen Testing (PhenoSys)

Rats were subcutaneously implanted with RFID transponders under isoflurane anaesthesia and maintained at ∼90% free-feeding body weight. Pre-training stages (Habituation through Punish Incorrect) shaped touchscreen-directed behaviour using a three-window mask [40], with sessions capped at 30 min or 30 trials and a 1 h inter-session interval (see **Supplementary Methods**). For pairwise discrimination (PD) and reversal learning series (R1 or R2), a two-window mask was used. The PD criterion was >80% accuracy across 2 sessions (2 × 30 positive trials) within one day; the same criterion applied to RL. Rats were capped at 20 RL sessions to prevent overtraining. Four experimental cohorts were tested: ABA-naïve and ABA-exposed rats receiving drug one day prior to either PD or R1.

### Data Analysis

Statistical analyses were performed in GraphPad Prism 9.5.1 (p<0.05; trends at p<0.10). Tests included two-tailed unpaired t-tests, one- and two-way ANOVA with Bonferroni, Dunnett, or Sidak post hoc corrections, and mixed-effects models, selected based on data type and group structure. Full statistical details are provided in **Supplementary Tables 2 to 6**.

## Results

### Under food restriction, psilocybin improved reversal learning performance with an apparent temporal delay compared to prior *ad libitum* fed cohort

Standard operant-level calorie restriction altered the temporal pattern through which psilocybin enhanced reversal learning. Using the FED3 paradigm ([8] **Fig 1A**), accuracy improved over time (*p* < .001; **Fig 1B**) and psilocybin-treated rats reached performance criterion earlier, with improved accuracy evident on reversal day 2 (*p* < .05) compared to day 3 in saline controls (*p* < .001; **Fig 1C**). This was accompanied by increased pellets won (*p* < .01; day 2 psilocybin > saline, *p* < .05; **Fig 1D**) and target response rates (*p* < .001; day 2 psilocybin > saline, *p* < .05; **Fig 1E**), without changes in overall response vigour (*p* = .174; **Fig 1F**). Non-target responding decreased over time (*p* < .001), with saline-treated rats trending toward greater perseverative responding on day 3 (*p* = .0843; **Fig 1G**), suggesting poorer inhibitory learning relative to psilocybin-treated animals. For *ad libitum* comparisons (**Fig 1H–J**), we performed additional analyses on our previously published dataset [8], collected on identical FED3 hardware, schedules, and in the same facility and light-cycle, to reveal that psilocybin-treated, *ad libitum* fed rats displayed steeper learning curves than their food-restricted counterparts (*p* < .01; **Fig 1I**), with no such difference in saline-treated animals (*p*=.106; **Fig 1J**), indicating that calorie restriction selectively attenuated psilocybin’s capacity to enhance the rate of new learning. This was not explained by motivational differences, as sucrose intake was equivalent across groups (*p*=.306; **Fig 1K**). Psilocybin transiently reduced home-cage chow intake in the 0–2 h (*p* < .001) and 2–5 h (*p* < .001) windows post-administration, that dissipated prior to operant testing times (*p*=.548; **Fig 1L**). Consistent with a ventral striatal contribution to these learning differences, psilocybin elicited greater NAc cFos+ expression (**Fig 1M**) across both nutritional states with food-restricted conditions exacerbating this impact both in terms of cFos+ area (*p* < .01: **Fig 1N**), and total cFos+ cell counts (*p* < .05; **Fig 1O**).

**Fig. 1.**
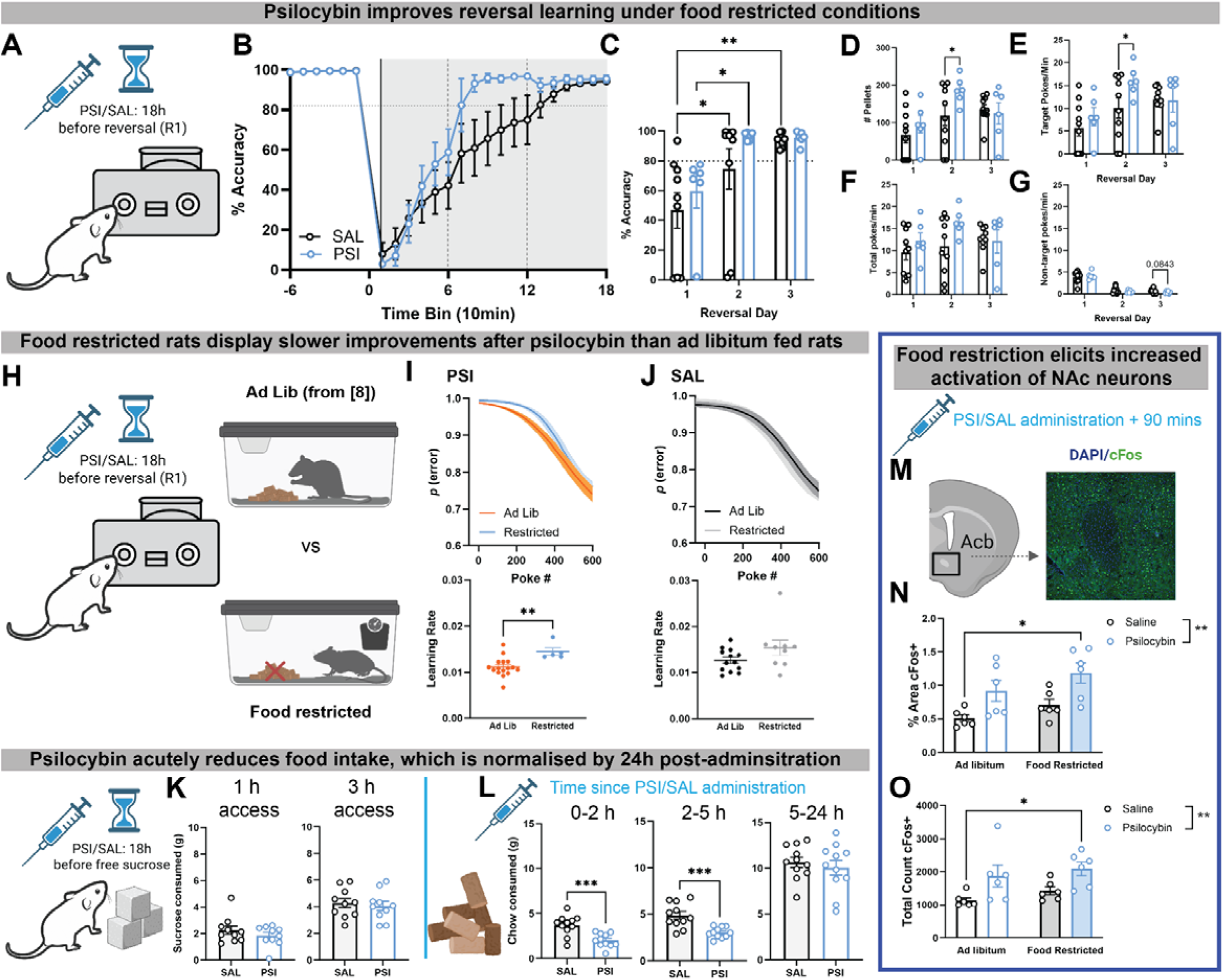
Standard calorie restriction for operant procedures alters post-acute psilocybin effects on new learning. Schematic of the FED3 operant device and between-session reversal learning timeline (**A**). Saline (SAL)- and psilocybin (PSI)-treated rats were tested across three daily reversal sessions (3 h each); response accuracy is plotted in 10 min time bins across days, with individual traces shown for visualisation only **(B**). A main effect of reversal day was observed (F^(2,25)^ = 15.9, *p* < .001), with no significant effect of drug or interaction. Post-hoc analyses revealed that psilocybin improved accuracy from reversal day 1 to day 2 (*p* < .05), enabling psilocybin-treated rats to reach performance criterion earlier than saline-treated rats, who reached criterion only on day 3 (*p* < .001; **C**). A main effect of reversal day was also observed for pellets won (F^(2,27)^ = 8.176, *p* < .01), with psilocybin-treated rats earning more pellets than saline controls on reversal day 2 (*p* < .05; **D**). Target poke response vigour (pokes/min) similarly increased across reversal days (F^(2,27)^ = 9.284, *p* < .001), with psilocybin-treated rats showing greater target response rates on reversal day 2 relative to saline (*p* < .05; **E**), indicating enhanced reinforcement learning. Total poke rate per minute did not differ significantly between treatment groups across days (**F**). Non-target response rates decreased over time (F^(1,21)^ = 110.3, *p* < .001), and while no effect of psilocybin was observed on day 2 non-target responding, saline-treated rats trended towards higher non-target response rates on day 3 (*p* = .0843; **G**), suggesting slower extinction of incorrect responding. Schematic depicting the fully fed (*ad libitum*) versus food-restricted (calorie-restricted) FED3 operant design (**H**). Psilocybin-treated rats maintained on *ad libitum* chow displayed an enhanced learning rate relative to their food-restricted counterparts (*t*(19) = 3.235, *p* < .01), while no corresponding difference in learning rate was observed between feeding conditions in saline-treated rats (**I**). No significant differences between fed and fasted animals were observed in the midpoint of the learning curve for either treatment (**J**). Post-acute sucrose consumption, assessed 24 h after administration consistent with FED3 testing parameters, was not affected by prior psilocybin treatment at either 1 h or 3 h test durations (**K**). Psilocybin reduced acute chow intake in the 0–2 h (*t*(20) = 4.094, *p* < .001) and 2–5 h (*t*(20) = 4.218, *p* < .001) post-administration windows, with no significant differences observed in the 5–24 h window (**L**). Psilocybin elicited greater cFos+ expression in the nucleus accumbens (**M**) across both *ad libitum* and food-restricted conditions (F^(1,5)^ = 22.32, *p* < .01), with post-hoc analyses confirming significantly greater expression in psilocybin-treated food-restricted rats compared to *ad libitum* saline-treated rats (*p* < .05; **N**). This result was replicated using the total count of cFos+ cells (F^(1,5)^ = 15.36, *p* < .05; **O**). Grouped data show mean ± SEM with individual data points. \**p* < .05, \*\**p* < .01, \*\*\**p* < .001. SAL saline, PSI psilocybin, FED feeding experimentation device. For complete details of statistical tests see **Supplementary Table 2.**

### Psilocybin amplifies NAc dopamine dynamics during probabilistic reversal learning

A serial probabilistic reversal learning task revealed sustained effects of psilocybin on cognitive flexibility over seven post-administration days while leveraging the dynamic nature of within-session reversals. Event-aligned responses were aggregated across the seven post-administration days per subject to improve precision in this small photometry cohort (n=4/group). Behavioural measures trended toward psilocybin-enhanced performance, including increased trial completion (*p* = .052; **Fig 2A**), successful reversals (*p* = .0802; **Fig 2B**), and overall wins (*p* = .0759; **Fig 2C**), without distinct differences in win-stay or lose-shift decision-making strategies or total losses (all *p’s* >.05; **Fig 2D-F**). Real-time NAc dopamine monitoring revealed that psilocybin significantly amplified dopamine transients time-locked to expected wins (*p* < .001; **Fig 2G,H**), expected losses (*p* < .001; **Fig 2I,J**), and unexpected wins at the non-target port (*p* < .01; **Fig 2K,L**), whereas unexpected losses produced robust dopamine dips regardless of treatment (*p*=.1727; **Fig 2M,N**), suggesting a ceiling effect. Computational model comparison (using BIC) revealed the Rescorla-Wagner model provided the better fit in all 9 rats (mean rat-level BIC: RW = 1171.35, AI = 1306.42; mean rat-level AIC: RW = 1097.43, AI = 1183.22; refer to **Supplementary Results**). Rescorla-Wager modelling of post-administration learning performance [46] revealed psilocybin-specific shifts in learning rate (α: *d* ≈ +1.1), prior value weighting (V_₀_: *d* ≈ −1.16), and exploration (β: *d* ≈ −0.6), indicating enhanced feedback-driven updating and reduced reliance on prior expectations (**Fig 2O,P**) effects absent under control conditions.

**Fig. 2.**
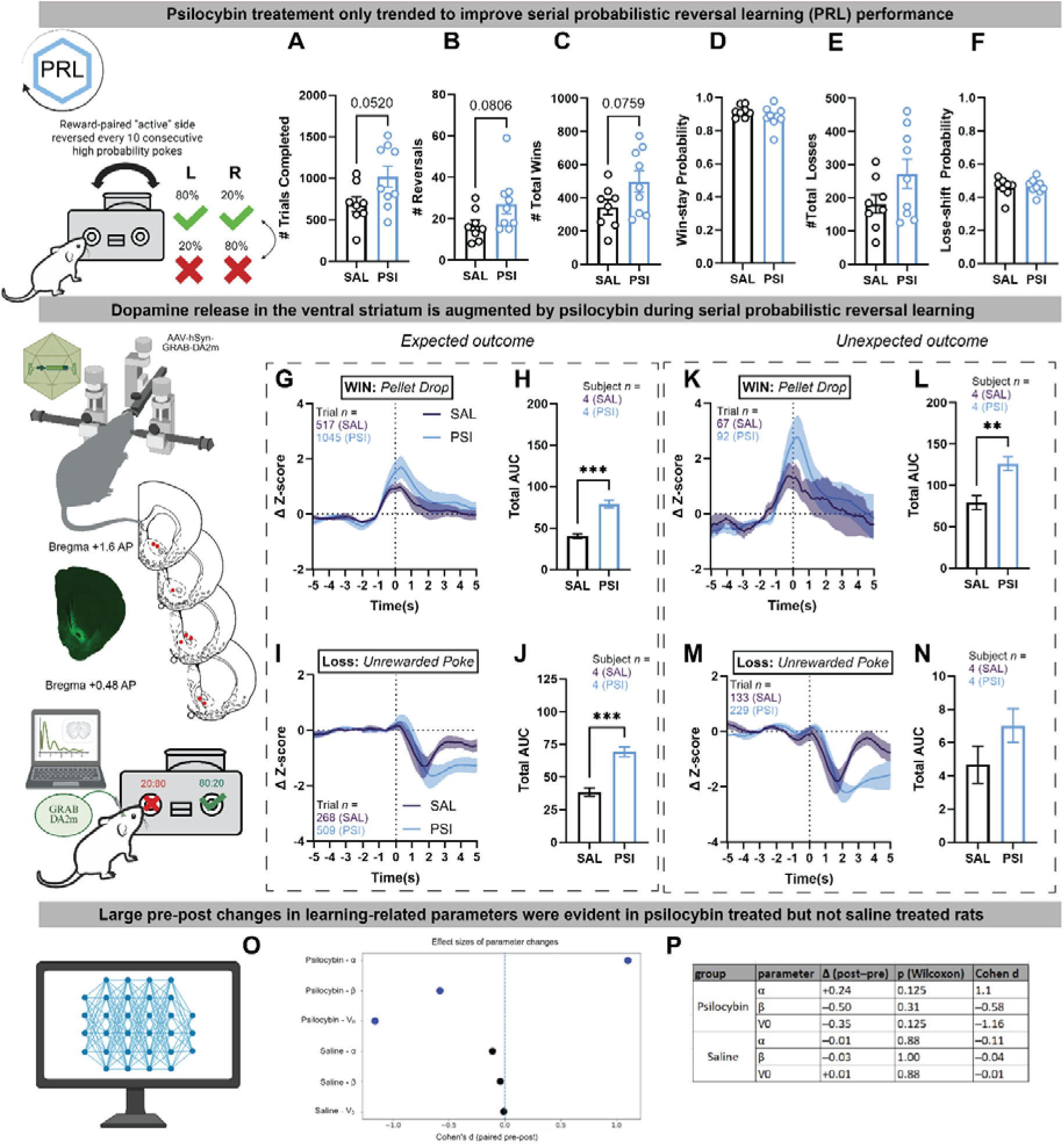
Dopamine reward signals in the ventral striatum are enhanced by psilocybin post-acutely during serial probabilistic reversal learning. Schematic of the serial probabilistic reversal learning task with within-session reversals, allowing tracking of post-acute psilocybin effects over one week of testing (**A**). Psilocybin trended towards increasing the total number of trials completed across seven days of testing (*t*(15) = 2.111, *p* = .0520; **B**), and similarly trended towards increasing the number of within-session reversals completed (*t*(15) = 1.874, *p* = .0802; **C**) and the total number of wins (*t*(15) = 1.907, *p* = .0759; **D**). No significant differences were observed in the probability of win-stay responding (**E**), the total number of losses (**F**), or the probability of lose-shift responding (**G**). Fibre photometry recordings of dopamine (DA) binding in the ventral striatum (VStr) were time-locked to task outcomes. Psilocybin significantly increased the area under the curve (AUC) of the DA transient in response to expected rewarded outcomes (target port pellet delivery; *t*(6) = 7.302, *p* < .001; **G-H**). DA transients were also enhanced for expected losses (unrewarded non-target pokes), with the AUC significantly greater in psilocybin-treated rats (*t*(6) = 6.182, *p* < .001; **I-J**). For unexpected rewarded outcomes (pellet delivery at the non-target port), psilocybin again increased DA transients, reflected in a significantly greater AUC (*t*(6) = 3.994, *p* < .01; **K-L**). Psilocybin sustained the DA dip in response to unexpected losses (unrewarded pokes at the target port), however this did not reach statistical significance in AUC, potentially reflecting a ceiling effect whereby the unexpected nature of these outcomes produces robust dips in both groups (**M-N**). Computational modelling using the Rescorla–Wagner framework revealed directional pre–post changes in learning-related parameters specifically following psilocybin treatment. Psilocybin produced a large positive effect on the learning rate (α: *d* ≈ +1.1), consistent with enhanced feedback-driven updating, whereas saline produced negligible change. A large negative shift in the prior value parameter (V₀: *d* ≈ −1.16) was observed after psilocybin only, suggesting down-weighting of prior expectations. A moderate negative effect on the exploration/noise parameter (β: *d* ≈ −0.6) was also observed with psilocybin, indicating a possible increase in choice stochasticity, though this effect was more variable (**O**). Parameter estimates are summarised in (**P**). Grouped data show mean ± SEM with individual data points. \*\**p* < .01, \*\*\**p* < .001. SAL saline, PSI psilocybin, DA dopamine, VStr ventral striatum, AUC area under the curve. For complete details of statistical tests see **Supplementary Table 3.**

### Psilocybin effects on discrimination accuracy and reversal depend on prior learning experience

The touchscreen-based visual discrimination and reversal learning task dissected stage-specific effects of psilocybin on goal-directed learning (**Fig 3A**). When administered prior to initial visual pairwise discrimination (PD), psilocybin reduced days (*p* < .01; **Fig 3B**) and sessions (*p* < .05; **Fig 3C**) to reversal (R1) criterion, without affecting trials to criterion (*p*=.3864; **Fig 3D**), suggesting that the total action-outcome experience required was unchanged. Psilocybin treatment reduced session completion time during initial discrimination (*p* = .0506; **Fig 3E**) and increased the overall proportion of correct responses (*p* < .0001; *p* < .05; **Fig 3F**), without effects on incorrect responding or omissions (*p*=.7400; **Fig 3G**; *p*=.1174; **Fig 3H**). Examining the first seven PD sessions individually confirmed that psilocybin significantly increased discrimination accuracy (Session: *p* < .0001; Drug: *p* < .01; **Fig 3i**) and correspondingly decreased inaccuracy (Session: *p* < .0001; Drug: *p* < .05; **Fig 3j**), indicating a robust effect of psilocybin across the early learning period. When administered prior to R1 instead (**Fig 3K**), psilocybin had no effect on reversal acquisition (all *p’s* >.05; **Fig 3L-N**), suggesting that reversal learning rate improvements only emerge when psilocybin is administered prior to initial discrimination learning. While a trend toward improved session completion time at the serial reversal learning stage (R2) was observed (*p* = .0820; **Fig 3O**), psilocybin significantly impaired accuracy (Training Stage: *p* < .05; Drug: *p* < .05; **Fig 3P**), without effects on incorrect responses (**Fig 3Q**), omissions (*p*=.1859; **Fig 3R**), or accuracy at R1 (all *p’s* >.05; **Fig 3s,t**). Together, these findings indicate that psilocybin differentially modulates initial goal-directed and subsequent flexible learning depending on training stage, with benefits contingent on administration prior to initial discrimination.

**Fig. 3.**
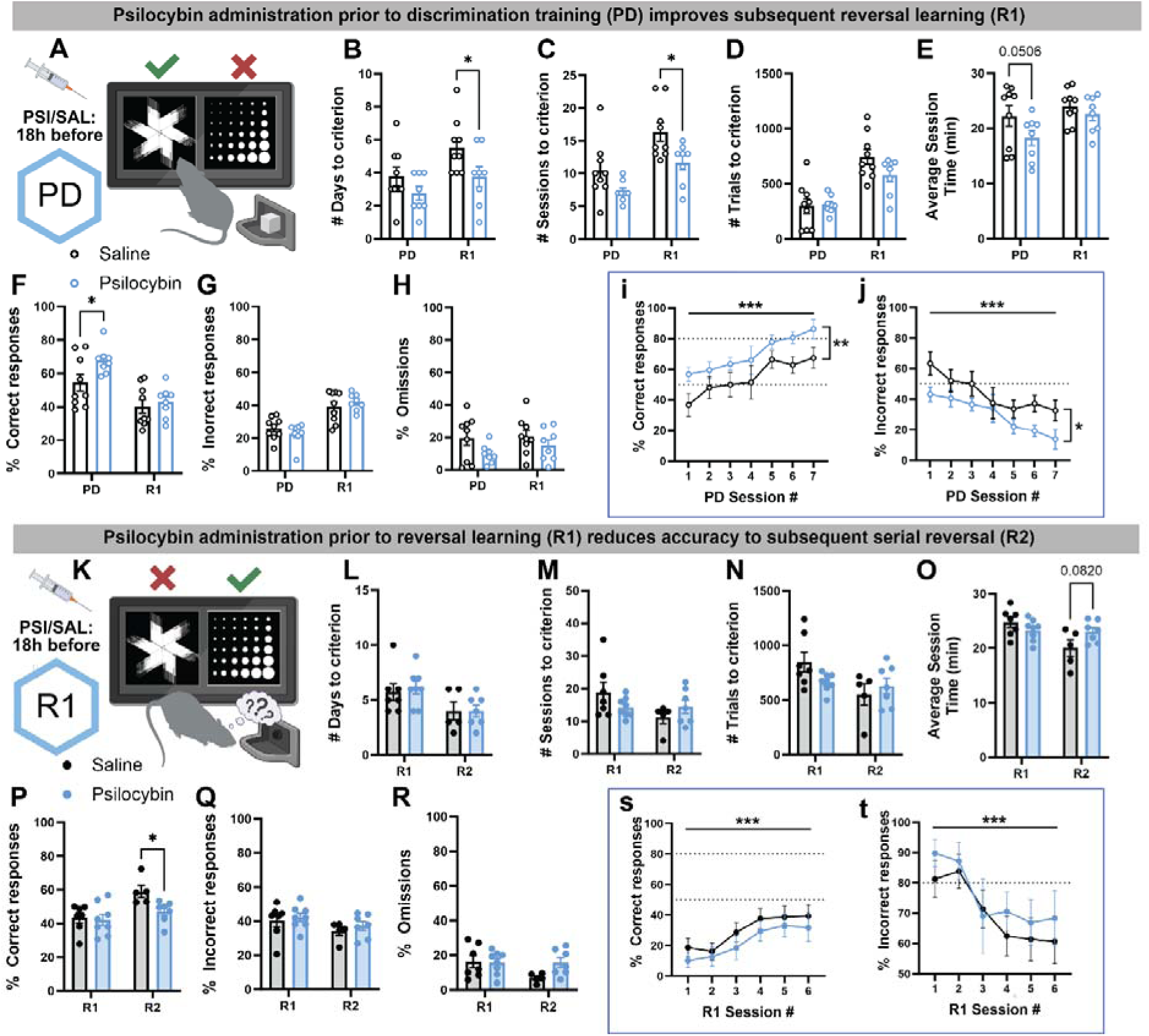
Psilocybin differentially improves goal-directed (PD) and flexible learning (R1, R2) in wild-type rats. Schematic depicting the PhenoSys operant design in which psilocybin or saline was administered prior to initial goal-directed learning (positive discrimination; PD), followed by serial reversal learning (R1, R2; **A**). Psilocybin reduced the number of days required to reach criterion at R1, with main effects of training stage (F^(1,9)^ = 10.02, *p* < .01) and drug treatment (F^(1,9)^ = 6.078, *p* < .05), and post-hoc analysis confirming improved R1 acquisition in psilocybin-treated rats (*p* < .05; **B**). This was mirrored by a reduction in sessions to criterion at R1 (Training Stage: F^(1,9)^ = 10.02, *p* < .01; Drug: F^(1,9)^ = 6.078, *p* < .05; **C**). No significant effect of psilocybin was observed on the number of trials to criterion (**D**), suggesting that while psilocybin accelerated acquisition over time, the total volume of action–outcome experience required was not altered. Psilocybin trended towards reducing average session completion time during PD only (*p* = .0506; **E**). Psilocybin improved the proportion of correct responses during PD (Training Stage: F^(1,9)^ = 112.9, *p* < .0001), with post-hoc analyses confirming greater accuracy in psilocybin-treated rats during PD (*p* < .05) but not at R1 (**F**). No significant effects were observed for incorrect responses (**G**) or omissions (**H**). Examination of the first seven PD sessions (up to the point the first animal reached criterion) revealed that psilocybin significantly increased accuracy (Session: F^(6,54)^ = 10.95, *p* < .0001; Drug: F^(1,9)^ = 10.71, *p* < .01; **i**) and correspondingly reduced inaccuracy (Session: F^(6,54)^ = 12.12, *p* < .0001; Drug: F^(1,9)^ = 6.719, *p* < .05; **j**) during this early learning phase. In a separate cohort, psilocybin or saline was administered prior to the first reversal (R1), with subsequent testing through serial reversal (R2; **K**). No significant effects of drug treatment were observed on days to criterion (**L**), sessions to criterion (**M**), or trials to criterion at R1 (**N**), indicating that improvements in reversal acquisition rate are specific to drug administration prior to initial discrimination learning. A trend towards reduced session completion time was observed at R2 in psilocybin-treated rats (*p* = .0820; **O**). Strikingly, psilocybin administered prior to R1 significantly improved accuracy at the subsequent serial reversal (R2; Training Stage: F^(1,7)^ = 8.351, *p* < .05; Drug: F^(1,7)^ = 7.59, *p* < .05; post-hoc *p* < .05; **P**). No significant effects of drug were observed for incorrect responses (**Q**) or omissions at R2 (**R**), and psilocybin had no effect on accuracy (**s**) or inaccuracy (**t**) at R1 itself. Grouped data show mean ± SEM with individual data points. \**p* < .05, \*\**p* < .01, \*\*\**p* < .001. SAL saline, PSI psilocybin, PD positive discrimination, R1 first reversal, R2 second reversal. For complete details of statistical tests see **Supplementary Table 4.**

### Prior activity-based anorexia exposure disrupts psilocybin-enhanced discrimination accuracy and reversal learning

To determine whether ABA exposure (**Fig 4A1**) altered psilocybin’s pro-cognitive effects, we administered psilocybin prior to PD in ABA-exposed rats, with equivalent weight loss, food intake and running wheel activity between treatment groups (all *p’s* >.05; **Fig 4A2**). ABA exposure had no effect on time to reach PD criterion (**Fig 4B-D**), but reduced discrimination accuracy (*p* < .05; **Fig 4E**); psilocybin improved accuracy in control animals (*p* < .01) but not ABA-exposed rats. Incorrect responses, omissions, and session completion times were unaffected (all *p’s* >.05; **Fig 4F-H**). During subsequent reversal learning, ABA exposure did not significantly alter days or sessions to reach R1 criterion (all *p’s* >.05; **Fig 4 I-K**) but markedly increased the number of trials required (*p* < .0001; **Fig 4L**) regardless of treatment. ABA exposure also reduced R1 accuracy (*p* < .0001; **Fig 4M**), which was not driven by changes in incorrect responses (*p =* .0720; **Fig 4N**) but by increased omissions (*p* < .01; **Fig 3O**), suggesting disengagement from the task rather than altered response selection. However, while ABA exposure prolonged R1 session completion time (*p* < .01); **Fig 4P**, psilocybin treatment reduced session completion time overall (*p* = .0518), suggesting a preservation of task engagement, in line with our previous work [47], despite the broader learning impairments induced by ABA exposure.

**Fig. 4.**
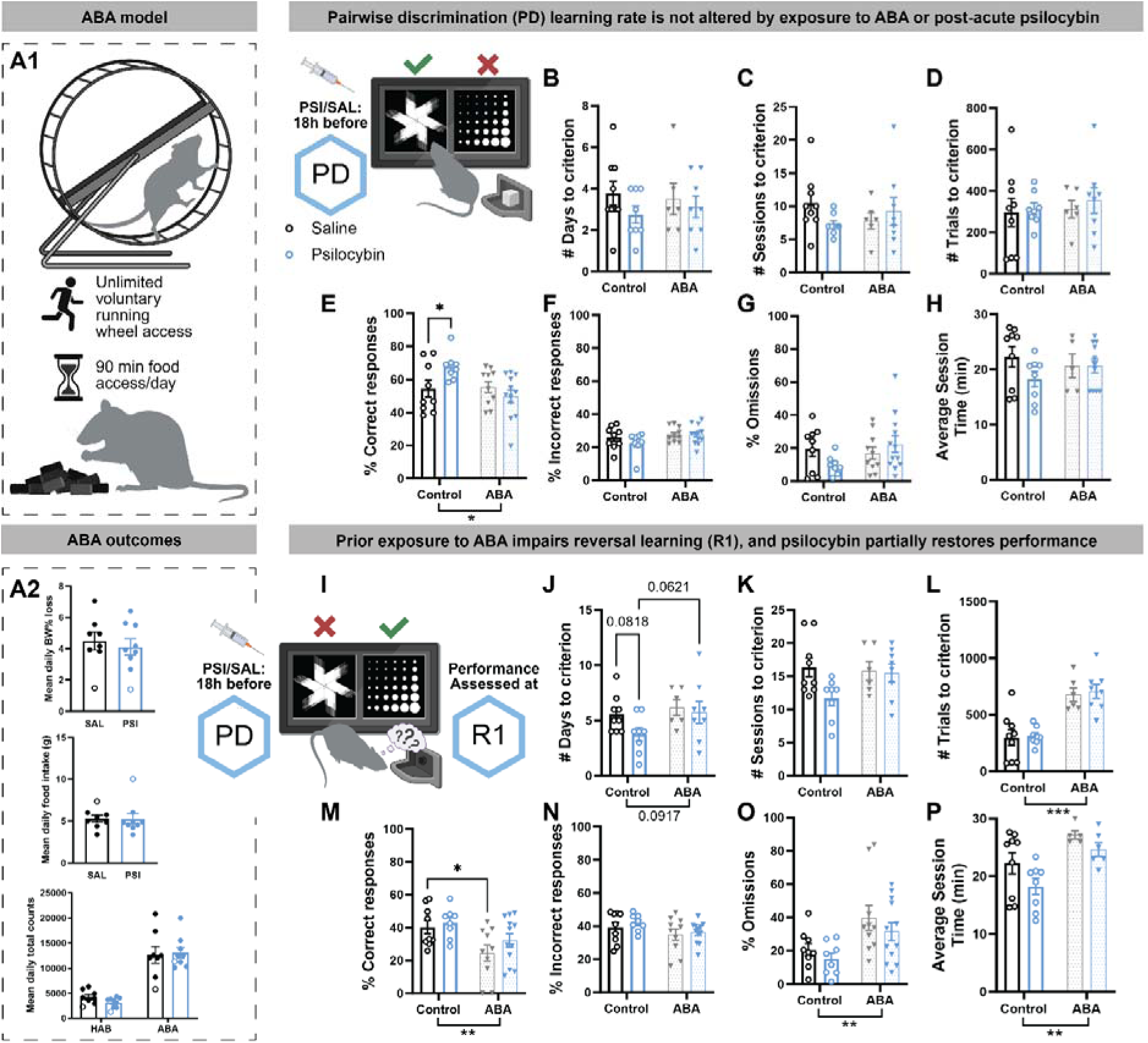
Psilocybin administered prior to PD does not rescue visual discrimination or reversal learning deficits observed in ABA. Schematic of the experimental design in which activity-based anorexia (ABA) or control rats received psilocybin or saline prior to commencement of positive discrimination (PD) training in the PhenoSys operant system (**A**). No significant effects of drug treatment or ABA exposure were observed on the number of days (**B**), sessions (**C**), or trials (**D**) required to reach PD criterion. ABA exposure reduced overall PD accuracy (ABA Exposure: F^(1,35)^ = 4.482, *p* < .05), and within control animals, psilocybin improved the proportion of correct responses relative to saline (*p* < .01), suggesting that ABA exposure disrupted psilocybin’s ability to enhance accurate responding during initial discrimination (**E**). No significant effects were observed for incorrect responses (**F**), omissions (**G**), or average session completion time (**H**). The effect of prior PD psilocybin administration on subsequent reversal learning (R1) in ABA rats was then examined (**I**). ABA exposure trended towards increasing the days required to reach R1 criterion (ABA Exposure: F^(1,27)^ = 3.058, *p* = .0917), and while psilocybin trended towards improving this measure (*p* = .0818), ABA-exposed psilocybin-treated rats trended towards poorer reversal acquisition relative to non-ABA psilocybin-treated rats (*p* = .0621; **J**). No significant effects were observed for sessions to criterion (**K**). ABA exposure significantly increased the number of trials required to reach R1 criterion (ABA Exposure: F^(1,27)^ = 42.97, *p* < .0001), with post-hoc analyses confirming that ABA-exposed rats, regardless of treatment, required more action–outcome experience to achieve criterion (both *p* < .001; **L**). ABA exposure also significantly reduced R1 accuracy (ABA Exposure: F^(1,27)^ = 42.97, *p* < .0001), with significant impairment in saline-treated ABA rats (*p* < .05) and a trend in psilocybin-treated ABA rats (*p* = .0825; **M**). ABA exposure reduced the proportion of incorrect responses (ABA Exposure: F^(1,35)^ = 3.442, *p* < .01; **N**), seemingly driven by an increased proportion of omissions (ABA Exposure: F^(1,35)^ = 9.858, *p* < .01; **O**). ABA exposure prolonged R1 session completion time (ABA Exposure: F^(1,24)^ = 12.58, *p* < .01), with psilocybin trending towards reducing this in all animals (Drug: F^(1,24)^ = 4.189, *p* = .0518; **P**). Grouped data show mean ± SEM with individual data points. \**p* < .05, \*\**p* < .01, \*\*\**p* < .001, \*\*\*\**p* < .0001. SAL saline, PSI psilocybin, ABA activity-based anorexia, PD positive discrimination, R1 first reversal. For complete details of statistical tests see **Supplementary Table 5.**

### Psilocybin administered prior to reversal learning produces opposing effects on accuracy and task engagement in ABA-exposed rats

When psilocybin was administered prior to R1 in ABA-exposed rats **(Fig 5A,B**), no effects of ABA exposure or drug treatment were observed on reversal learning acquisition across days, sessions, or trials to criterion (all *p’s* >.05; **Fig 5C-E**), nor on the proportion of correct responses, incorrect responses, or omissions (all *p’s* >.05; **Fig 5F-H**). Counterintuitively, ABA exposure reduced session completion time during reversal learning (*p* < .05; **Fig 5I**), an effect driven specifically by saline-treated animals (*p* < .05). During subsequent serial reversal learning (R2; **Fig 5J**), a significant interaction between ABA exposure and drug treatment was observed for accuracy (*p* < .05; **Fig 5K**), with psilocybin worsening accuracy specifically in control rats (*p* < .05), without significant effects on incorrect responses (*p =* .3388; **Fig 5L**) or omissions (*p =* .1149; **Fig 5M**). A parallel interaction was also observed for average session completion time (*p* < .05; **Fig 5N**), with psilocybin specifically reducing session completion time in ABA-exposed rats (*p* < .05). The opposing direction of effects is consistent with a speed–accuracy trade-off; however, in the absence of trial-wise latency analyses this interpretation remains provisional.

**Fig. 5.**
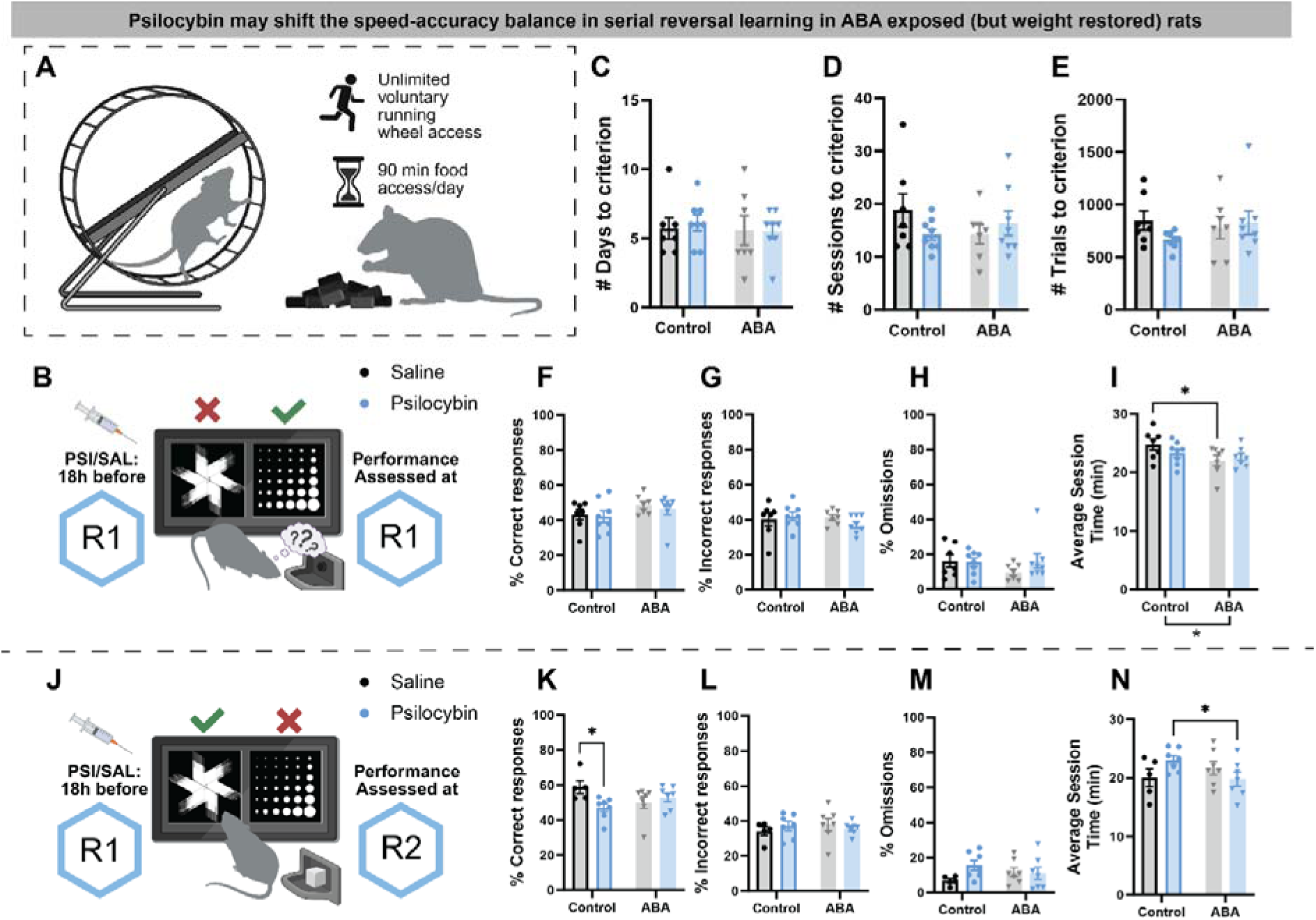
Psilocybin may shift the speed–accuracy balance in serial reversal learning performance in ABA. Schematic of ABA induction (**A**) followed by psilocybin or saline administration prior to first reversal learning (R1; **B**). No significant effects of ABA exposure or drug treatment were observed on days to criterion (**C**), sessions to criterion (**D**), or trials to criterion (**E**) at R1. No significant effects were found for the proportion of correct (**F**), incorrect (**G**), or omitted (**H**) responses. Counterintuitively, ABA exposure significantly reduced R1 session completion time (ABA Exposure: F^(1,26)^ = 4.938, *p* < .05), an effect driven specifically by saline-treated ABA rats (*p* < .05; **I**). Schematic of ABA induction followed by drug administration prior to serial reversal learning (R2; **J**). A significant interaction between ABA exposure and drug treatment was observed for R2 accuracy (Interaction: F^(1,22)^ = 6.032, *p* < .05), with psilocybin reducing accuracy in control but not ABA rats (*p* < .05; **K**). No significant main effects or interactions were observed for incorrect responses (**L**) or omissions (**M**). A significant interaction between ABA exposure and drug treatment was also observed for average R2 session completion time (Interaction: F^(1,22)^ = 4.605, *p* < .05), with psilocybin reducing session completion time across both control and ABA groups relative to controls (*p* < .05; **N**). Together, the observation that psilocybin reduced accuracy in control animals while simultaneously reducing session completion time in both groups suggests a possible shift in the speed–accuracy trade-off following psilocybin administration prior to serial reversal learning. Whether this represents a true trade-off or increased response vigour warrants further investigation. Grouped data show mean ± SEM with individual data points. \**p* < .05. SAL saline, PSI psilocybin, ABA activity-based anorexia, R1 first reversal, R2 second reversal. For complete details of statistical tests see **Supplementary Table 6.**

## Discussion

Here, we show that the cognitive improvements elicited by psilocybin are supported, in part, by an amplification of nucleus accumbens (NAc) dopamine transients, which may operate alongside broader serotonergic signalling and neuroplastic changes to accelerate learning rates. However, these cognitive benefits are gated by current metabolic state and prior experience with pathological weight loss. Ultimately, these results indicate that metabolic disruption and stress may alter the neurobiological substrates required for psilocybin-induced plasticity, suggesting a need to interpret therapeutic outcomes in the context of baseline physiological state (**Fig. 6**).

**Fig. 6.**
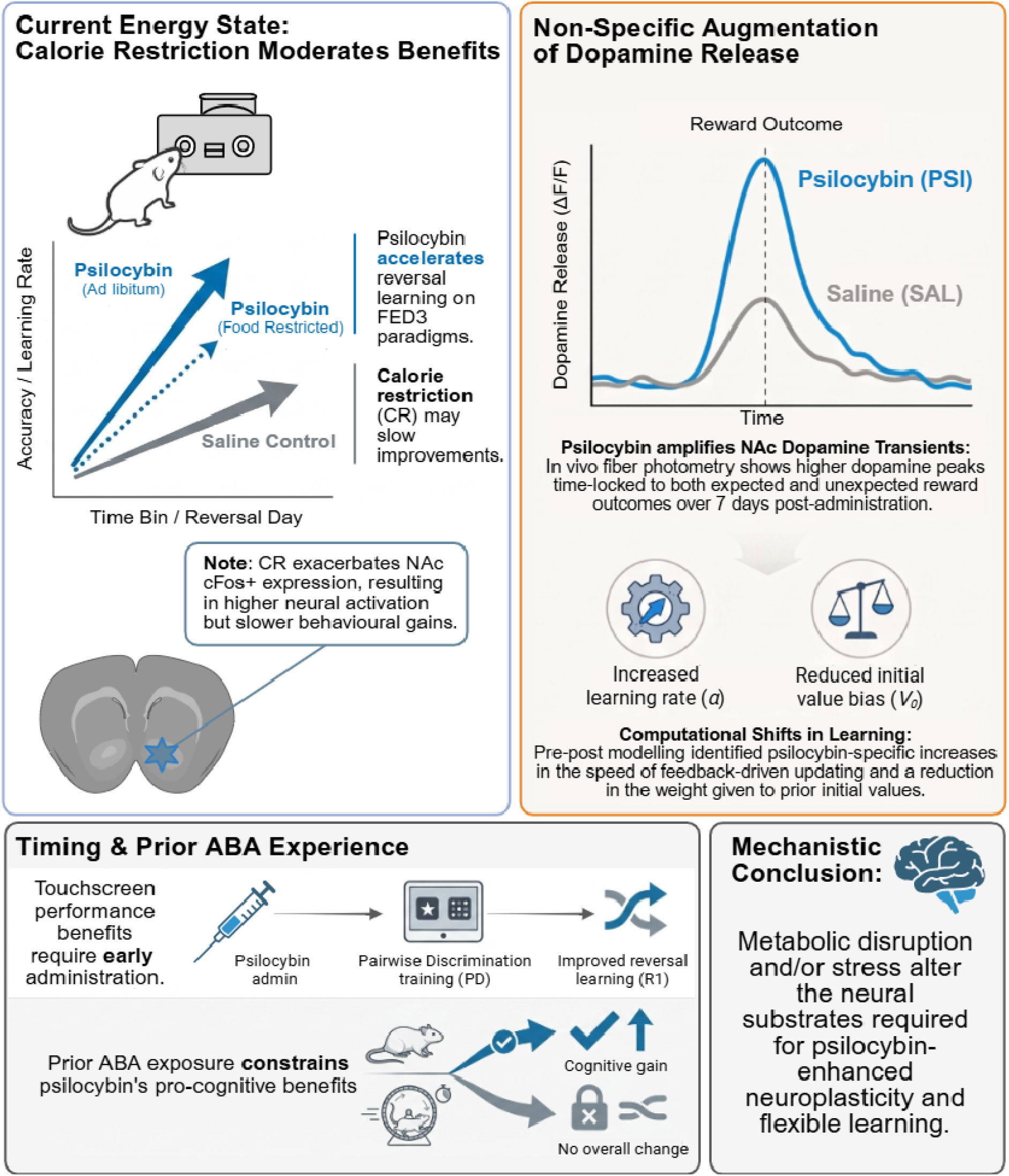
Synthesis of experimental findings. We show that 1) calorie restriction may moderate the benefits of psilocybin on reversal learning and exacerbates activation of NAc neurons; 2) improvements in learning after psilocybin are facilitated by augmentation of dopamine release to both expected and unexpected reward outcomes; 3) timing of administration within the learned task structure is important for subsequent learning and; 4) prior exposure to activity-based anorexia (ABA) constrains benefits for cognitive performance. Taken together, these data suggest that metabolic disruption and/or stress alter the neura substrates required for psilocybin-enhanced neuroplasticity and flexible learning.

### Context-dependent effects of psilocybin on cognitive flexibility: task structure, food availability, and test environment

Our findings demonstrate that psilocybin’s capacity to enhance cognitive flexibility is shaped by the conditions under which learning occurs. Across three paradigms, improvements in flexible learning were consistently modulated by nutritional state, prior ABA exposure, and task structure, aligning with the view that psychedelics act as modulators of neural plasticity whose effects are gated by environmental and experiential factors [10, 48, 49]. Calorie restriction delayed psilocybin’s capacity to accelerate new learning, which was unlikely to reflect motivational differences given equivalent sucrose intake across groups. Food restriction alters serotonergic tone, 5-HT2A receptor expression, and dopaminergic sensitivity in ways that may constrain the neural substrate available for psilocybin-induced plasticity [50], and the exacerbated cFos+ expression observed under food restriction in the NAc suggests that caloric deprivation amplifies striatal activation without proportional behavioural benefit.

Divergent findings across touchscreen learning stages may reflect the influence of prior learning experience, whereby differences in task demands and, consequently, recruitment of neural circuitry [51] alter the effects of psilocybin. When comparing stage-specific performance at the point of administration, the lack of effect on reversal performance, alongside distinct effects on initial discrimination, suggests that these improvements reflect genuine cognitive enhancement rather than overall changes in motivation or motor function. Unlike spatially anchored FED3 contingencies, touchscreen visual discrimination requires stimulus-specific memory encoding that may be more vulnerable to the memory-impairing effects of psilocybin reported elsewhere [11]. If psilocybin can simultaneously impair memory encoding while enhancing cognitive flexibility [7, 8], then the opposing effects we observe across reversal stages may reflect precisely this dissociation. In support of this interpretation is evidence that psilocybin robustly enhances fear extinction in mice when administered acutely prior to testing, whereas administered prior to fear learning or immediately after extinction yields no change in behaviour [48]. Together, these findings underscore the importance of task architecture and testing context when interpreting studies of psilocybin-enhanced cognition and brain function, with implications for how clinical trials are structured with respect to set and setting.

### Dopaminergic contributions to psilocybin-enhanced cognition: beyond a purely serotonergic mechanism

To our knowledge, these data provide the first *in vivo* evidence consistent with psilocybin-driven modulation of NAc dopamine transients during probabilistic reversal learning. While 5-HT2A receptor agonism dominates mechanistic accounts of psilocybin’s therapeutic effects [52], no neurochemical system operates in isolation [15]. Psilocybin broadly amplified NAc dopamine transients time-locked to expected and unexpected outcomes, with this amplification emerging throughout the seven days beyond the acute psychedelic state, implying enduring reorganisation of reward prediction error signalling rather than a transient pharmacological effect. Our findings align with the elevated GRAB-DA signal observed in rats administered the psychedelic 5-HT2A/C agonist DOI [53], in which both expected and unexpected rewards elicited an increase in NAc dopamine release in a Pavlovian task, although this prior study investigated only acute drug effects.

This broad amplification of reinforcement signalling suggests that psilocybin facilitates adaptive behaviour by strengthening the neural encoding of actionable feedback signals over time. The absence of effects of psilocybin on unexpected losses likely relates to a ceiling effect whereby the inherently surprising nature of these outcomes already maximally engages prediction error mechanisms regardless of treatment. These findings complement recent transcriptomic evidence showing upregulated plasticity genes in intratelencephalic neurons at 72 hours post-psilocybin administration [54], suggesting that the sustained dopaminergic effects observed here may reflect enduring neuroplastic changes that fundamentally alter how striatal circuits process reinforcement. A critical question remains about whether these effects stem from direct dopaminergic action or represent downstream consequences of initial serotonergic activation, owing to the well-described and potent agonist action of psilocybin at 5-HT receptor subtypes. Altered serotonin transmission has been linked to neuronal responses to unpredicted or surprising events, and compared to dopamine neuron activity is specifically important for inhibiting perseverative responding [55]. Thus, it is likely that the changes in dopamine and learning shown here are combined effects of both transmitter systems.

Furthermore, these enduring psilocybin-induced changes align with the cortico-striatal cascade of psychedelic effects in which 5-HT2A receptor activation on cortical pyramidal neurons drives glutamate release that subsequently modulates striatal dopamine activity [56]. Computational modelling reinforced this by revealing that psilocybin produced large shifts from baseline in learning rate and prior value weighting consistent with strengthened feedback-driven updating, in line with enhanced dopamine prediction error signalling. The disorder-specificity of these dopaminergic effects warrants emphasis. In AN, where NAc hypodopaminergia reduces reward sensitivity and impairs adaptive goal pursuit [57], amplification of dopamine prediction error signals may restore motivational salience of food-related outcomes. However, augmented dopamine signalling in AN may also drive food avoidance in AN [36] and dopamine system aberrations may differ based on illness stage [37]. Additionally, in substance use disorders characterised by dopaminergic hypersensitivity [58], the same mechanism may be counterproductive, highlighting that psilocybin’s dopaminergic actions are likely dependent on disorder state, as well as context. Future studies using operant paradigms in familiar versus novel contexts would help clarify whether environmental novelty during testing contributes to the dopamine dynamics observed here.

### Activity-based anorexia exposure constrains psilocybin’s cognitive benefits: implications for therapeutic outcomes

ABA-exposed rats failed to show psilocybin-enhanced reversal learning, challenging a “deficit-remediation” account whereby psychedelics preferentially benefit individuals with the greatest baseline impairment. The notion that psychedelic efficacy may scale with baseline impairment is conceptually consistent with other neuromodulatory interventions, such as methylphenidate in attention-deficit/hyperactivity disorder, where individuals with lower striatal dopamine, as well as those with higher trait impulsivity, show greater improvements in reversal learning performance following treatment [59]. Overall, our findings suggest that while psilocybin may enhance cognitive flexibility in intact systems, it may not improve cognitive performance in an animal model with a known impairment measured using the same operant task design [40]. This is partially consistent with the clinical literature: Both clinical trials evaluating psilocybin in women with AN published to date [3, 38] reported heterogeneous psilocybin responses in women with AN, with those carrying the most entrenched illness trajectories *least* likely to benefit [62]. However, it is important to note that exposure to the ABA model is a stressful experience [60] that can have enduring effects on behaviour [61], which may interact with psilocybin in ways that have not been examined here. In humans, psilocybin-induced increases in neural flexibility are not consistently accompanied by behavioural improvements across pathological states [4], further highlighting an important boundary condition on the relationship between psilocybin’s cognitive effects and baseline neurobiological state.

It is plausible that ABA-associated metabolic or neurobiological changes interfere with psilocybin’s mechanisms of action relevant to cognitive flexibility. Our previous work demonstrated that ABA exacerbates acute reductions in cortical 5-HT2A receptor expression elicited by psilocybin [8], which would be expected to attenuate the cortico-striatal glutamatergic drive proposed to underlie psilocybin-enhanced dopamine signalling. While direct comparisons are limited, both clinical and preclinical datasets suggest that illness chronicity, current nutritional state and associated neural adaptations may moderate psilocybin response. The variability in response to psilocybin in both humans and preclinical models carries an important practical implication: studies evaluating psilocybin’s cognitive effects likely require substantially larger sample sizes than those typically employed to detect treatment effects and identify responder subgroups. In light of this, the present study highlights some trend-level observations that may emerge as important features of psilocybin effects if adequately powered to detect response stratified subgroups, echoing the recent call for larger sample sizes in clinical trials of psilocybin in eating disorder populations [66]. Stratifying animals by severity of ABA-induced cognitive impairment, rather than treating exposed animals as homogeneous, may reveal whether a threshold of neurobiological disruption exists below which psilocybin retains cognitive benefits [67].

### Future directions: bridging neurochemical synergy and clinical translation in anorexia nervosa

The present findings raise the question of whether neurochemical tone at the time of treatment moderates psilocybin’s cognitive effects. Given our prior observation that ABA exposure exacerbated reductions in cortical 5-HT2A receptor expression [8], this may be one mechanism contributing to the constrained response to psilocybin, although the present study suggests that analogous changes in striatal dopamine receptor availability may impose a similar ceiling. Positron emission tomography (PET) imaging with specific radioligands could provide a non-invasive index of baseline neurochemical function in patients prior to treatment [68], and whether pre-treatment dopaminergic tone predicts therapeutic outcome, independently of or synergistically with other brain-based mechanisms is a tractable clinical question. This is particularly pressing given that neither dopaminergic nor serotonergic pharmacotherapies alone have demonstrated robust efficacy in AN [69], therefore psilocybin’s therapeutic logic may lie in its capacity to engage both systems simultaneously. A multi-modal biomarker approach that moves beyond single-system characterisation may be valuable in this context. Blood-based profiling of the tryptophan-serotonin and kynurenine pathways offers a tractable, minimally invasive index of serotonergic substrate availability that could be deployed alongside PET imaging to capture both peripheral neurochemical state and central receptor availability within the same individual. The convergence of these measures into an integrated predictive framework represents the logical next step for the field [70]. Rather than asking whether psilocybin works in AN, the field is now positioned to ask a more precise and clinically actionable question: for which neurochemical profile does it work best, and can that profile be identified before treatment begins? Answering this will require the kind of bench-to-bedside translational architecture in which preclinical biomarker signatures are prospectively validated in parallel clinical trials. Such an approach will transform response variability from an obstacle into a mechanistic window into the neurobiology of psychedelic-assisted recovery.

## Acknowledgements

We thank Monash undergraduate students Amelia Trice and Meagan Lee for contributing to data collection for some of these studies, and Dr Beth Fisher (Monash) and A/Prof Ryan Smith (Laureate Institute for Brain Research) for assistance with computational modelling frameworks. We acknowledge the USONA Institute Investigational Drug Supply Program for providing the psilocybin used in these studies and A/Prof Alexxai Kravitz, Washington University in St. Louis for input into operant task design and operations. We acknowledge the use of the Monash Microimaging Facility and Biorender for aspects of the figures. We declare that generative AI (Claude.ai) was used to scan for redundant text in the final drafting stages of the initial submission to reduce word count.

## References

1. Goodwin, G.M., et al., Single-Dose Psilocybin for a Treatment-Resistant Episode of Major Depression. N Engl J Med, 2022. 387(18): p. 1637–1648.

2. Meshkat, S., et al., Efficacy and safety of psilocybin for the treatment of substance use disorders: A systematic review. Neuroscience & Biobehavioral Reviews, 2025. 173: p. 106163.

3. Peck, S.K., et al., Psilocybin therapy for females with anorexia nervosa: a phase 1, open-label feasibility study. Nature Medicine, 2023. 29(8): p. 1947–1953.

4. Doss, M.K., et al., Psilocybin therapy increases cognitive and neural flexibility in patients with major depressive disorder. Transl Psychiatry, 2021. 11(1): p. 574.

5. Sloshower, J., et al., Psychological flexibility as a mechanism of change in psilocybin-assisted therapy for major depression: results from an exploratory placebo-controlled trial. Scientific Reports, 2024. 14(1): p. 8833.

6. Meshkat, S., et al., Impact of psilocybin on cognitive function: A systematic review. Psychiatry Clin Neurosci, 2024. 78(12): p. 744–764.

7. Torrado Pacheco, A., Olson, R.J., Garza, G., and Moghaddam, B., Acute psilocybin enhances cognitive flexibility in rats. Neuropsychopharmacology, 2023. 48(7): p. 1011–1020.

8. Conn, K., et al., Psilocybin restrains activity-based anorexia in female rats by enhancing cognitive flexibility: contributions from 5-HT1A and 5-HT2A receptor mechanisms. Mol Psychiatry, 2024. 29(10): p. 3291–3304.

9. Shadani, S., Greaves, E., Andrews, Z.B., and Foldi, C.J., Psilocybin exerts differential effects on social behavior and inflammation in mice in contexts of activity-based anorexia. Psychedelics, 2026. **-**1(aop): p. 1–12.

10. Rijsketic, D.R., et al., UNRAVELing the synergistic effects of psilocybin and environment on brain-wide immediate early gene expression in mice. Neuropsychopharmacology, 2023. 48(12): p. 1798–1807.

11. Rambousek, L., Palenicek, T., Vales, K., and Stuchlik, A., The effect of psilocin on memory acquisition, retrieval, and consolidation in the rat. Front Behav Neurosci, 2014. 8: p. 180.

12. Gregersen, A.T., Whelan, T., Golden, C., Blackmore, T., and Palner, M., Computational Analysis of Psilocybin Effects on Three-Choice Touchscreen Reversal Learning in Rats: A Pilot Study. Psychedelic Medicine, 2026: p. 28314425251386879.

13. Carter, O.L., et al., Using Psilocybin to Investigate the Relationship between Attention, Working Memory, and the Serotonin 1A and 2A Receptors. Journal of Cognitive Neuroscience, 2005. 17(10): p. 1497–1508.

14. Cameron, L.P., et al., Beyond the 5-HT(2A) Receptor: Classic and Nonclassic Targets in Psychedelic Drug Action. J Neurosci, 2023. 43(45): p. 7472–7482.

15. McCoy, K., Reed, F., Conn, K., and Foldi, C.J., Separate or inseparable? Serotonin and dopamine system interactions may underlie the therapeutic potential of psilocybin for anorexia nervosa. Physiol Behav, 2025. 298: p. 114957.

16. Aghajanian, G.K. and Marek, G.J., Serotonin, via 5-HT2A receptors, increases EPSCs in layer V pyramidal cells of prefrontal cortex by an asynchronous mode of glutamate release. Brain Research, 1999. 825(1): p. 161–171.

17. Welch, A.C., et al., Dopamine D2 receptor overexpression in the nucleus accumbens core induces robust weight loss during scheduled fasting selectively in female mice. Mol Psychiatry, 2021. 26(8): p. 3765–3777.

18. Foldi, C.J., Milton, L.K. and Oldfield, B.J., The Role of Mesolimbic Reward Neurocircuitry in Prevention and Rescue of the Activity-Based Anorexia (ABA) Phenotype in Rats. Neuropsychopharmacology, 2017. 42(12): p. 2292–2300.

19. Beeler, J.A., et al., Vulnerable and Resilient Phenotypes in a Mouse Model of Anorexia Nervosa. Biol Psychiatry, 2021. 90(12): p. 829–842.

20. Klenotich, S.J., Ho, E.V., McMurray, M.S., Server, C.H., and Dulawa, S.C., Dopamine D2/3 receptor antagonism reduces activity-based anorexia. Translational Psychiatry, 2015. 5(8): p. e613–e613.

21. Limbrick-Oldfield, E.H., van Holst, R.J. and Clark, L., Fronto-striatal dysregulation in drug addiction and pathological gambling: Consistent inconsistencies? NeuroImage: Clinical, 2013. 2: p. 385–393.

22. Celone, K.A., Thompson-Brenner, H., Ross, R.S., Pratt, E.M., and Stern, C.E., An fMRI investigation of the fronto-striatal learning system in women who exhibit eating disorder behaviors. NeuroImage, 2011. 56(3): p. 1749–1757.

23. Uniacke, B., et al., Resting-state connectivity within and across neural circuits in anorexia nervosa. Brain Behav, 2019. 9(1): p. e01205.

24. Zastrow, A., et al., Neural correlates of impaired cognitive-behavioral flexibility in anorexia nervosa. Am J Psychiatry, 2009. 166(5): p. 608–16.

25. Watabe-Uchida, M., Eshel, N. and Uchida, N., Neural Circuitry of Reward Prediction Error. Annu Rev Neurosci, 2017. 40: p. 373–394.

26. Deng, Y., Song, D., Ni, J., Qing, H., and Quan, Z., Reward prediction error in learning-related behaviors. Frontiers in Neuroscience, 2023. **Volume** 17 - 2023.

27. Vollenweider, F.X., Vontobel, P., Hell, D., and Leenders, K.L., 5-HT modulation of dopamine release in basal ganglia in psilocybin-induced psychosis in man--a PET study with [11C]raclopride. Neuropsychopharmacology, 1999. 20(5): p. 424–33.

28. Sakashita, Y., et al., Effect of psilocin on extracellular dopamine and serotonin levels in the mesoaccumbens and mesocortical pathway in awake rats. Biol Pharm Bull, 2015. 38(1): p. 134–8.

29. Wierenga, C.E., et al., Hunger does not motivate reward in women remitted from anorexia nervosa. Biol Psychiatry, 2015. 77(7): p. 642–52.

30. Wierenga, C.E., Brown, C.S. and Reilly, E.E., Reinforcement Learning and Decision Making in Anorexia Nervosa. Curr Psychiatry Rep, 2025. 27(12): p. 711–722.

31. Heshmati, M. and Russo, S.J., Anhedonia and the brain reward circuitry in depression. Curr Behav Neurosci Rep, 2015. 2(3): p. 146–153.

32. Pothos, E.N., Creese, I. and Hoebel, B.G., Restricted eating with weight loss selectively decreases extracellular dopamine in the nucleus accumbens and alters dopamine response to amphetamine, morphine, and food intake. J Neurosci, 1995. 15(10): p. 6640–50.

33. Kaye, W.H., Frank, G.K.W. and McConaha, C., Altered Dopamine Activity after Recovery from Restricting-Type Anorexia Nervosa. Neuropsychopharmacology, 1999. 21(4): p. 503–506.

34. Carr, K.D., Tsimberg, Y., Berman, Y., and Yamamoto, N., Evidence of increased dopamine receptor signaling in food-restricted rats. Neuroscience, 2003. 119(4): p. 1157–67.

35. Thanos, P.K., Michaelides, M., Piyis, Y.K., Wang, G.J., and Volkow, N.D., Food restriction markedly increases dopamine D2 receptor (D2R) in a rat model of obesity as assessed with in-vivo muPET imaging ([11C] raclopride) and in-vitro ([3H] spiperone) autoradiography. Synapse, 2008. 62(1): p. 50–61.

36. Frank, G.K.W. From Desire to Dread—A Neurocircuitry Based Model for Food Avoidance in Anorexia Nervosa. Journal of Clinical Medicine, 2021. 10, DOI: 10.3390/jcm10112228.

37. Beeler, J.A. and Burghardt, N.S., The Rise and Fall of Dopamine: A Two-Stage Model of the Development and Entrenchment of Anorexia Nervosa. Frontiers in Psychiatry, 2022. **Volume** 12 **-** 2021.

38. Douglass, H.M., et al., Psilocybin therapy for adult females with anorexia nervosa: pilot study. The British Journal of Psychiatry, 2026: p. 1–9.

39. Cora, M.C., Kooistra, L. and Travlos, G., Vaginal Cytology of the Laboratory Rat and Mouse: Review and Criteria for the Staging of the Estrous Cycle Using Stained Vaginal Smears. Toxicol Pathol, 2015. 43(6): p. 776–93.

40. Huang, K., et al., Rapid, automated, and experimenter-free touchscreen testing reveals reciprocal interactions between cognitive flexibility and activity-based anorexia in female rats. eLife, 2023. 12: p. e84961.

41. Milton, L.K., et al., Suppression of Corticostriatal Circuit Activity Improves Cognitive Flexibility and Prevents Body Weight Loss in Activity-Based Anorexia in Rats. Biological Psychiatry, 2021. 90(12): p. 819–828.

42. Schneider, C.A., Rasband, W.S. and Eliceiri, K.W., NIH Image to ImageJ: 25 years of image analysis. Nat Methods, 2012. 9(7): p. 671–5.

43. Matikainen-Ankney, B.A., et al., An open-source device for measuring food intake and operant behavior in rodent home-cages. Elife, 2021. 10.

44. Lopes, G. and Monteiro, P., New Open-Source Tools: Using Bonsai for Behavioral Tracking and Closed-Loop Experiments. Front Behav Neurosci, 2021. 15: p. 647640.

45. He, Q., Liu, J.L., Eschapasse, L., Beveridge, E.H., and Brown, T.I., A comparison of reinforcement learning models of human spatial navigation. Scientific Reports, 2022. 12(1): p. 13923.

46. Pearce, J.M. and Bouton, M.E., Theories of associative learning in animals. Annu Rev Psychol, 2001. 52: p. 111–39.

47. Fisher, E.L., et al., Psilocybin increases optimistic engagement over time: computational modelling of behaviour in rats. Transl Psychiatry, 2024. 14(1): p. 394.

48. Woodburn, S.C., Levitt, C.M., Koester, A.M., and Kwan, A.C., Psilocybin Facilitates Fear Extinction: Importance of Dose, Context, and Serotonin Receptors. ACS Chem Neurosci, 2024. 15(16): p. 3034–3043.

49. Fadahunsi, N., et al., Acute and long-term effects of psilocybin on energy balance and feeding behavior in mice. Transl Psychiatry, 2022. 12(1): p. 330.

50. Portero-Tresserra, M., et al., Caloric restriction modulates the monoaminergic system and metabolic hormones in aged rats. Scientific Reports, 2020. 10(1): p. 19299.

51. Ghosh, S. and Zador, A.M., Corticostriatal Plasticity Established by Initial Learning Persists after Behavioral Reversal. eneuro, 2021. 8(2): p. ENEURO.0209-20.2021.

52. Drewko, A.J., Habets, R.L.P. and Brunt, T.M., Above the threshold, beyond the trip: the role of the 5-HT(2A) receptor in psychedelic-induced neuroplasticity and antidepressant effects. Mol Psychiatry, 2025. 30(12): p. 5926–5937.

53. Martin, D.A., Delgado, A.M. and Calu, D.J., Effects of psychedelic, DOI, on nucleus accumbens dopamine signaling to predictable rewards and cues in rats. Neuropsychopharmacology, 2024. 49(12): p. 1925–1933.

54. Liao, C., et al., Single-nucleus transcriptomics reveals time-dependent and cell-type-specific effects of psilocybin on gene expression. bioRxiv, 2025.

55. Matias, S., Lottem, E., Dugué, G.P., and Mainen, Z.F., Activity patterns of serotonin neurons underlying cognitive flexibility. eLife, 2017. 6: p. e20552.

56. Olson, D.E., Biochemical Mechanisms Underlying Psychedelic-Induced Neuroplasticity. Biochemistry, 2022. 61(3): p. 127–136.

57. Avena, N.M. and Bocarsly, M.E., Dysregulation of brain reward systems in eating disorders: neurochemical information from animal models of binge eating, bulimia nervosa, and anorexia nervosa. Neuropharmacology, 2012. 63(1): p. 87–96.

58. Boileau, I., et al., Conditioned dopamine release in humans: a positron emission tomography [11C]raclopride study with amphetamine. J Neurosci, 2007. 27(15): p. 3998–4003.

59. Clatworthy, P.L., et al., Dopamine release in dissociable striatal subregions predicts the different effects of oral methylphenidate on reversal learning and spatial working memory. J Neurosci, 2009. 29(15): p. 4690–6.

60. Burden, V.R., Douglas White, B., Dean, R.G., and Martin, R.J., Activity of the Hypothalamic-Pituitary-Adrenal Axis is Elevated in Rats with Activity-Based Anorexia123. The Journal of Nutrition, 1993. 123(7): p. 1217–1225.

61. Kinzig, K.P. and Hargrave, S.L., Adolescent activity-based anorexia increases anxiety-like behavior in adulthood. Physiol Behav, 2010. 101(2): p. 269–76.

62. Peck, S.K., et al., Psychedelic treatment for anorexia nervosa: A first-hand view of how psilocybin treatment did and did not help. Psychedelics, 2024. 1(1): p. 15–18.

63. Breton, J., et al., Characterizing the metabolic perturbations induced by activity-based anorexia in the C57Bl/6 mouse using (1)H NMR spectroscopy. Clin Nutr, 2020. 39(8): p. 2428–2434.

64. Prochazkova, P., et al., Microbiome and metabolic disruption in acute vs. severe and enduring anorexia nervosa. npj Biofilms and Microbiomes, 2025. 11(1): p. 217.

65. Foldi, C.J., Taking better advantage of the activity-based anorexia model. Trends Mol Med, 2024. 30(4): p. 330–338.

66. Bevione, F., Lacidogna, M.C., Lavalle, R., Abbate Daga, G., and Preti, A., Psilocybin in the treatment of eating disorders: a systematic review of the literature and registered clinical trials. Eat Weight Disord, 2025. 30(1): p. 58.

67. Mattay, V.S., et al., Effects of Dextroamphetamine on Cognitive Performance and Cortical Activation. NeuroImage, 2000. 12(3): p. 268–275.

68. Manza, P., et al., Neural basis for individual differences in the attention-enhancing effects of methylphenidate. Proceedings of the National Academy of Sciences, 2025. 122(13): p. e2423785122.

69. Frank, G.K.W., Pharmacotherapeutic strategies for the treatment of anorexia nervosa – too much for one drug? Expert Opinion on Pharmacotherapy, 2020. 21(9): p. 1045–1058.

70. Foldi, C.J. and Griffiths, K.R., Examining the biological causes of eating disorders to inform treatment strategies. Nat Rev Neurosci, 2025.

