## Supplementary for "Appetite for change: How nutritional state and ventral striatal dopamine shape psilocybin effects on cognitive flexibility"

### Text summary of Supplementary Materials

Contained in this document is all supplementary materials to accompany the manuscript entitled “Appetite for change: how nutritional state and ventral striatal dopamine shape psilocybin effects on cognitive flexibility” by Conn K, *et al* (2026).

It includes additional methodological information, animal allocation and exclusions, modelling methods and results and full details of all statistical analyses in tables.

### Supplementary Methods

#### Animals and Housing

All animals were obtained from the Monash Animal Research Platform (MARP; Clayton, VIC, Australia). Young female Sprague Dawley rats were used in these studies, based on their unique susceptibility to activity-based anorexia (ABA) and the translational relevance of assessing this group to understand the therapeutically relevant effects of psilocybin for anorexia nervosa. Rats were group-housed and acclimated to the 12 h light/dark cycle (lights off at 1100 h) for 7 days in a temperature (22-24°C) and humidity (30-50%) controlled room before experiments commenced and had ad libitum access to water and standard laboratory chow (Barastoc, Australia) unless undergoing food restriction procedures. Because aspects of reinforcement learning and ABA (i.e. wheel running and food intake) are known to fluctuate with the oestrous cycle in female rats [1, 2], a pair of male rats were housed in all experimental rooms at least 7 days prior to experimentation in order to facilitate synchronisation of cycling, known as the Whitten Effect [3]. For the ABA experiments, modelling began at 6 to 7 weeks of age as per Conn et al., 2024, for all feeding and operant testing experiments, rats were aged 10-12 weeks of age at the start of training.

For all behavioural experiments except for those requiring PhenoSys testing, animals were housed in specialised individual cages for the duration of testing (26 cm W x 21 cm H 47.5 cm D). For Phenosys testing, animals were housed in groups of 5 to 8 in specialised cages (26 cm × 34 cm × 55 cm) to allow for RFID implanted sorted of animals into the automated testing chamber as per [4]. For experiments conducted requiring food restriction, animals went under standard operant testing procedures where food provided daily was titrated to ensure that rats remained within 90-95% of their baseline body weight. All experimental procedures were conducted in accordance with the Australian Code for the care and use of animals for scientific purposes and approved by the Monash Animal Resource Platform Ethics Committee (ERM 36800).

### Supplementary Table 1. Animal numbers and experimental allocation

| **Experiment** | **Training** | **Subject (N)** | **Drug admin** | **Post-acute testing** |
| --- | --- | --- | --- | --- |
| Baseline feeding (chow) | N/A | N = 20 (n=10 SAL; n = 10 PSI) | Home-cage chow removed 30min prior to admin | Chow returned to home-cage 30 min after admin, consumption measured 2, 5, & 24h later |
| Baseline feeding (sucrose) | Previous 24h chow consumption test | As above | At 24h post-admin, chow removed | Free access to sucrose began at admin, sucrose consumption measured at 1 & 3h |
| Baseline Feeding (cFos+ Analysis) | Ad libitum vs. food restriction for 7 days | N = 24  n=6 SAL-*ad lib;* n=6 SAL-food restricted; n=6 PSI-*ad lib;* n=6 PSI-food restricted | Drug admin 7-days after the start of the food-access manipulation | Rats were collected for immunohistochemical validation of cFos+ expression 18-24h post-admin |
| Between-sessions Reversal Learning | FR1 (3 Days)  FR1 (3 Days)  FR5 (3 Days) | N = 15 (n=9 SAL; n=6 PSI) | 18h prior to FR5 Reversal (as per [5]) | FR5 Reversal (3 Days) – All under food restricted conditions |
| Within-session Reversal Learning | FR1 both (1-week)  100:0 (2 weeks) – Reversal after 6 consecutive correct responses | N = 17  For behaviour: n=8 SAL; n=9 PSI  For fibre photometry of N=17, n=4 SAL; n=4 PSI were selected | 18h prior to the introduction of 80:20 probabilistic reward contingencies | Fibre photometry recordings conducted for 7 days of probabilistic reversal learning (80:2) post-admin.  Rats were collected for immunohistochemical validation of GRAB-DA expression at end of recordings. |
| ABA-naive Phenosys Testing | Training as per [4] | Admin prior to PD: N=8-9/treatment  Admin prior to R1:  N=7/treatment | Drug administered either the day before (~18h prior to) beginning PD or R1 | For admin prior to PD:  Testing at R1🡪 R2  For admin prior to R1:  Testing at R2🡪 R3 |
| ABA PhenoSys Testing | 1-week post ABA exposure - Training as per [4] | Admin prior to PD: N=8-9/treatment  Admin prior to R1:  N=7-8/treatment |  | For admin prior to PD:  Testing at R1🡪 R2  For admin prior to R1:  Testing at R2🡪 R3 |

#### Stereotaxic Surgery and Viral Preparation

Rats were anaesthetised using 5% isoflurane with a flow rate of 0.5 L/min and positioned in a head-fixed stereotaxic apparatus, with the plane of anaesthesia being maintained at 2.5% at 0.5 L/min. With the aid of microinjector, a glass micropipette (Drummond, #5-000-1001-X10, Birmingham, AL, USA) was inserted into the core of the NAc (A-P: +1.70; M-L: +/- 1.50; D-V: -6.2 from bregma), and 400nl of virus encoding the dopamine biosensor (pAAV-hsyn-GRAB_DA2m; Addgene, #140553, Watertown, MA, USA) was infused unilaterally into the area at a rate of 80 nl/min. An additional five-minute waiting time was used to allow for the diffusion of the virus before withdrawing the injector from the tissue. A Fibre optic cannula with black ceramic ferrule (core diameter of 200um; length of 7.5mm; numerical aperture of 0.37; ferrule size of 1.25mm; RWD Life Sciences) was then inserted into the same site. The sizes of the fibres were matched with that of the connecting patch cables (i.e., core = 200um, NA = 0.37; Doric Lenses, Québec, Canada) used during the photometry experiments. The ferrules were fixated into the skull using four stainless steel screws (length 4mm x width 1mm) implanted into each bone plate of the exposed skull. Finally, a solid head cap that covers the entire skull was created with light-cured dental cement (Gaenial Universal Flow; GC Australia Dental) to protect the exposed skull as well as holding the ferrule and screw heads in place. Surgeries were completed at least seven days prior to controlled room habituation to allow for adequate recovery.

#### Within-session Probabilistic Reversal Learning (FED) and In Vivo Fibre Photometry

We used a serial probabilistic reversal learning paradigm (PRL), which assesses subjects' ability to deploy the 'win-stay' and 'lose-shift' strategies, thereby assessing cognitive flexibility. During the testing process, rats were exposed to two unconditioned stimuli – an 'active' port with a high probability of being rewarded (i.e., 100% or 80%) and an 'inactive' port with a low probability of being rewarded (i.e., 0% or 20%). The orientation of the 'active' and 'inactive' ports was randomised at the start of each experiment. Subjects were required to guide their responses towards the 'active' port with the evidence for which action would lead to a reward collected across trials. Once the active port has been chosen for a designated number of times (in this case 10 consecutive high probability pokes), the reward contingencies were swapped without signalling the subject. Subjects were then required to apply the 'win-stay' and 'lose-shift' strategies again, flexibly adjust their behaviours in response to the new cue-outcome couples and select the port with the highest probability of being rewarded. This process was repeated throughout the 1h experimental cycle each day. Cognitive flexibility was quantified by the number of successful reversals and the probability of shifting to the opposite port after encountering a 'loss'.

For in vivo fiber photometry recordings, fluorescence signals were time-locked to FED3-coordinated nose-poke and reward delivery events and acquired using Bonsai software (Version 2.7.1[6]). Dual excitation LEDs (470 nm and 415 nm) were alternately pulsed at 40 Hz. The 470 nm signal corresponded to GRAB-DA sensor excitation and increased with dopamine binding, enabling real-time measurement of dopamine fluctuations in freely moving animals. The 415 nm signal served as an isosbestic control, providing a dopamine-independent reference. Behavioural analyses included n=8–9 animals per treatment group. Animals failing to meet performance criteria during deterministic reversal learning (≥3 reversals/session) were excluded a priori. Fibre photometry analyses were conducted on n=4 animals per group following post hoc exclusion for incorrect fibre placement or excessive signal noise. Animals were euthanised and brain tissue collected as described above. Coronal sections were imaged to verify GRAB-DA sensor expression and fibre placement within the NAc, and only animals with confirmed expression and correct targeting were included in analyses.

#### Pairwise Discrimination and Reversal Learning Touchscreen Testing (PhenoSys)

Rats were anaesthetized with 5% isoflurane and implanted subcutaneously with RFID transponders, with incision sites sealed using tissue adhesive. Thereafter, rats were maintained on daily feeding prior to the dark phase to sustain ~90% of free-feeding body weight throughout the experiment. A series of pre-training stages including Habituation, Initial Touch, Must Touch, Must Initiate, and Punish Incorrect were used to shape reward-based behaviors of the rats toward the touchscreen (as per Huang et al., 2023). A mask with three side-by-side windows was used in all pre-training stages. Rats were allowed to have multiple sessions of training per day, with a maximum duration of 30 min or 30 trials per session and a 1 hr time-out period between sessions. The pairwise discrimination (PD) and reversal learning (RL) task were used to assess cognitive flexibility in rats. A touchscreen mask with two side-by-side windows was used in the task. Rats were allowed to perform multiple sessions per day, as in pre-training stages, and were first required to discriminate between two stimulus images and associate touching one of the images with receiving the reward.

Rats were required to complete 2 sessions (2 × 30 positive trials) with accuracy >80% within one day to reach the progression criterion to reversal learning, in which the stimulus-reward association was reversed. The progression criterion in reversal learning remained the same as in pairwise discrimination. To assess flexible learning with or without prior exposure to ABA, each rat was restricted to a maximum of 20 sessions of reversal learning to prevent touchscreen overtraining. Once rats reached either the progression criterion (i.e. learned the task) or 20 sessions of reversal learning (i.e. did not learn the task), they were removed from experimentation. Four experiments were conducted using this training and testing paradigm: 1) wild-type rats administered drug prior to PD, 2) wild-type rats administered drug prior to R1, 3) ABA-exposed rats administered drug prior to PD and 4) ABA-exposed rats administered drug prior to R1.

#### Computational Modelling

*Reinforcement Learning Rescorla–Wagner model.*

The RL RW model included three free parameters: learning rate α, inverse temperature β, and initial value bias V_0_. On each trial t, the rat selected an action a_t_ ∈ {L,R}, corresponding to the left or right nose-poke port. Expected values for the two actions were initialised as:

$\mathbf{V}_{\mathbf{L}}\mathbf{(1)=}\mathbf{V}_{\mathbf{R}}\left( \mathbf{1} \right)\mathbf{=}\mathbf{V}_{\mathbf{0}}$

Choice probability was generated by a softmax decision rule over the action values:

$$\mathbf{p}\left( \mathbf{a}_{\mathbf{t}}\mathbf{=i} \right)\mathbf{=}\frac{\boldsymbol{exp(\beta}\mathbf{V}_{\mathbf{i}}\left( \mathbf{t} \right)\mathbf{)}}{\sum_{\mathbf{j}} \exp\left( \boldsymbol{\beta}\mathbf{V}_{\mathbf{j}}\left( \mathbf{t} \right) \right)}$$

where i,j ∈ {L,R}.
r_t_ was coded as 1 for reward and 0 for no reward in the present implementation. After the observed outcome (r_t_), the expected values (V) were updated using a standard reward prediction error:

$\mathbf{RP}\mathbf{E}_{\mathbf{t}}\mathbf{=}\mathbf{r}_{\mathbf{t}}\mathbf{-}\mathbf{V}_{\mathbf{at}}\left( \mathbf{t} \right)$

$\mathbf{V}_{\mathbf{at}}\left( \mathbf{t+1} \right)\mathbf{=}\mathbf{V}_{\mathbf{at}}\left( \mathbf{t} \right)\boldsymbol{+\alpha\cdot RP}\mathbf{E}_{\mathbf{t}}$

Where $V_{a}\left( t \right)$is value of the chosen action at trial t. Model fitting minimised the negative log-likelihood of the observed choices.

*Active Inference model.* In the AI model, beliefs about reward and loss outcomes under each action were encoded by Dirichlet concentration parameters collected in a likelihood matrix A. Initial beliefs were determined by a prior reward probability parameter pr, such that reward concentrations were initialised to pr×50 and loss concentrations to (1−pr)×50 for both actions. Outcome preferences were represented by a softmax-transformed preference vector with fixed reward value and fitted loss aversion LA:

$\mathbf{C=softmax}\left( \left[ \mathbf{2,-LA} \right] \right)$

For each action j, where j∈{L,R}, expected free energy combined an epistemic term and a pragmatic term:

$\mathbf{Q}_{\mathbf{j}}\mathbf{=-epistemi}\mathbf{c}_{\mathbf{j}}\mathbf{-pragmati}\mathbf{c}_{\mathbf{j}}$

where the epistemic term reflected expected information gain and the pragmatic term reflected expected log preference over outcomes. Collecting these into the vector Q=[Q_L_, Q_R_], action probabilities were generated using an inverse temperature parameter β:

$\mathbf{q}_{\mathbf{t}}\mathbf{=softmax}\left( \mathbf{-Q} \right)$

$\boldsymbol{\pi}_{\mathbf{t}}\mathbf{=softmax}\left( \boldsymbol{\beta}\log\mathbf{q}_{\mathbf{t}} \right)$

After each observed outcome, the Dirichlet concentration parameters for the chosen action were updated using separate decay terms for rewarded and unrewarded trials. Here, a_t_ denotes the Dirichlet concentration vector for the chosen action at trial t, and o_t_ denotes the observed outcome vector on that trial:

$\mathbf{a}_{\mathbf{t+1}}\mathbf{=}\left( \mathbf{1-}\boldsymbol{\alpha}_{\mathbf{decay}} \right)\mathbf{a}_{\mathbf{t}}\mathbf{+}\boldsymbol{\alpha}_{\mathbf{update}}\mathbf{o}_{\mathbf{t}}$

The update scaling was held fixed at 0.5 so that α_update_ = 0.5 [7]. The active inference parameters were estimated separately for each rat-session by minimising a variational-Laplace objective in transformed parameter space. In practice, optimisation was performed in Python using the L-BFGS-B algorithm. Because L-BFGS-B returns an approximate inverse Hessian, this was extracted and used as an approximate posterior covariance matrix for the fitted parameters. The fitted active inference parameters were action precision (β), loss aversion (LA), separate decay parameters for rewarded and unrewarded trials, and prior reward probability (pr).

#### Data Analysis

Statistical analyses were performed using GraphPad Prism 9.5.1 (GraphPad Software, San Diego, CA, USA). Statistical significance was set at *p* < 0.05, with *p* < .10 considered a trend though not significant. Analyses used were two-tailed unpaired t test, one-way and two-way analysis of variance (ANOVA) with Bonferroni’s, Dunnett’s or Sidak’s post hoc multiple comparisons, and a mixed-effects model, chosen appropriately considering the type of data, number of groups, and comparisons of interest. Full details of all statistical tests performed (including group composition) can be found in the **Supplementary Statistics Tables.**

### Supplementary Table 2. Statistical Summary of Experimental Outcomes for Figure 1 (over page)

| **Figure** | **Experimental Groups** | **Outcome Measure** | **Statistical Test** | **Main Statistics** | **Post-hoc Comparisons** |
| --- | --- | --- | --- | --- | --- |
| 1C | Food-restricted: SAL v PSI | % Accuracy by Day | Two-way ANOVA  [Factor 1 (Drug) x Factor 2 (Day)] | Reversal Day: F (2, 25) = 15.9, *P*<0.001; Drug: F(1, 13) = 1.211, *p*=.2911, ns. ; Interaction: F(2, 26) = 0.9164, *p*=.4125, ns. | *P*SI: Reversal Day 1 vs Day 2, *p* <.05; SAL: Reversal Day 1 vs Day 2, *p*<.05; SAL: Reversal Day 1 vs Day 2, *p*<.01. |
| 1D | Food-restricted: SAL v PSI | # Pellets | Two-way ANOVA  [Factor 1 (Drug) x Factor 2 (Day)] | Reversal Day: F (2, 27) = 8.176, *P*<.01; Drug: F(1, 14) = 1.329, *p*=.2863, ns. ; Interaction: F(2, 28) = 2.397, *p*=.1094, ns. | Reversal Day 2: SAL v *P*SI, *p*<.05 |
| 1E | Food-restricted: SAL v PSI | Target Pokes/min | Two-way ANOVA  [Factor 1 (Drug) x Factor 2 (Day)] | Reversal Day: F (2, 27) = 9.284, *p*<.001, *P*<.001; Drug: F(1, 14) = 1.652, *p*=.2196, ns. ; Interaction: F(2, 28) = 2.107, *p*=.1405, ns. | Reversal Day 2: SAL v *P*SI, *p*<.05 |
| 1F | Food-restricted: SAL v PSI | Total Pokes/min | Two-way ANOVA  [Factor 1 (Drug) x Factor 2 (Day)] | Reversal Day: F (2, 25) = 1.881, *P*=0.1736, ns; Drug: F(1, 13) = 1.621, *P*=0.2237, ns.; Interaction: F(2, 26) = 2,361, *P*=0.1129, ns. | None |
| 1G | Food-restricted: SAL v PSI | Non-target pokes/min | Two-way ANOVA  [Factor 1 (Drug) x Factor 2 (Day)/ | Reversal Day: F (1, 21) = 110.3, *P*<.001; Drug: F(1, 14) = 0.2720, *p*=.6101, ns. ; Interaction: F(2, 28) = 0.7728, *p*=.4713, ns. | Reversal Day 3: SAL v *P*SI, *p*=.0843 |
| 1I | Ad Lib v Fasted: PSI | Learning Rate | Independent  T-Test | t(19) =3.235, *p*<.01 | N/A |
| 1J | Ad Lib v Fasted: PSI | Learning Rate | Independent  T-Test | t(20) =1.695, *p*=.1055, ns. | N/A |
| 1K | Food intake (sated): SAL v PSI | 1h Sucrose consumption | Independent  T-Test | t(18)=1.053, *p*=.3064, ns. | N/A |
| 1K | Food intake (sated): SAL v PSI | 3h Sucrose consumption | Independent  T-Test | t(18)=0.4562, *p*=.6537, ns. | N/A |
| 1L | Food intake (sated): SAL v PSI | 0-2h Chow consumption | Independent  T-Test | t(20) =4.094, *p*<.001 | N/A |
| 1L | Food intake (sated): SAL v PSI | 2-5h Chow consumption | Independent  T-Test | t(20) =4.218, *p*<.001 | N/A |
| 1L | Food intake (sated): SAL v PSI | 5-24h Chow consumption | Independent  T-Test | t(20) =0.6119, *p*=.5475, ns. | N/A |
| 1N | Food-restricted: SAL v PSI | % Area cFos+ | Two-way ANOVA  [Factor 1 (Drug) x Factor 2 (Nutritional State)] | Food intake condition: F (1, 5) = 5.509, *p*<.05; Drug: F (1, 5) = 22.32, *p*<.01 ; Interaction: F(1, 5) = 2.107, *p*=.1405, ns. | Ad libitum:Saline vs. Food Restricted:*P*silocybin, *p*<.05 |
| 1O | Food-restricted: SAL v PSI | Total Count cFos+ | Two-way ANOVA  [Factor 1 (Drug) x Factor 2 (Nutritional State)] | Food intake condition: F (1, 5) = 1.128, *p*=.3368, ns.; Drug: F (1, 5) = 15.36, *p*<.05 ; Interaction: F(1, 5) = 0.01724, *p*=.9007, ns. | Ad libitum:Saline vs. Food Restricted:*P*silocybin, *p*<.05 |

### Supplementary Table 3. Statistical Summary of Experimental Outcomes for Figure 2

| **Figure #** | **Experimental Groups** | **Outcome Measure** | **Statistical Test** | **Main Statistics** |
| --- | --- | --- | --- | --- |
| 2A | Behaviour: SAL v PSI | # Trials Completed | Independent T-Test | t(15) =2.111, *p*=.0520, ns. |
| 2B | Behaviour: SAL v PSI | # Reversals | Independent T-Test | t(15) =1.874, *p*=.08016, ns. |
| 2C | Behaviour: SAL v PSI | # Total Wins | Independent T-Test | t(15) =1.907, *p*=.0759, ns. |
| 2D | Behaviour: SAL v PSI | Win-stay Probability | Independent T-Test | t(15) =1.293, *p*=.2155, ns. |
| 2E | Behaviour: SAL v PSI | # Total Losses | Independent T-Test | t(15) =1.686, *p*=.1124, ns. |
| 2F | Behaviour: SAL v PSI | Lose-shift Probability | Independent T-Test | t(15) =.2191, *p*=.8295, ns. |
| 2H | FiPho: SAL v PSI | Expected Win: Total AUC | Independent T-Test | t(6) =7.302, *p*<.001 |
| 2J | FiPho: SAL v PSI | Expected Loss:Total AUC | Independent T-Test | t(6) =6.182, *p*<.001 |
| 2L | FiPho: SAL v PSI | Unexpected Win:Total AUC | Independent T-Test | t(6) =3.994, *p*<.01 |
| 2N | FiPho: SAL v PSI | Unxpected Loss:Total AUC | Independent T-Test | t(6) =1.548, *p*=.1727, ns. |

### Supplementary Table 4. Statistical Summary of Experimental Outcomes for Figure 3

| **Figure #** | **Experimental Groups** | **Outcome Measure** | **Statistical Test** | **Main Statistics** | **Post-hoc Comparisons** |
| --- | --- | --- | --- | --- | --- |
| 3B | Pre-PD: WT SAL v PSI | # Days to criterion | Two-way ANOVA  [Factor 1 (Drug) x Factor 2 (Training Stage)] | Training Stage: F(1, 9) = 10.02, *p*<.01 ; Drug: F(1, 9) = 6.078, *p*<.05 ; Interaction: F(1, 3) = 1.218, *p*=.3540, ns. | R1: SAL-*P*SI; *p* <.05 |
| 3C | Pre-PD: WT SAL v PSI | # Sessions to criterion | Two-way ANOVA  [Factor 1 (Drug) x Factor 2 (Training Stage)] | Training Stage: F(1, 9) = 40.13, *p*<.0001 ; Drug: F(1, 9) = 9.013, *p*<.01 ; Interaction: F(1, 3) = 0.8731, *p*=.4190, ns. | R1: SAL-*P*SI; *p* <.05 |
| 3D | Pre-PD: WT SAL v PSI | # Trials to criterion | Two-way ANOVA  [Factor 1 (Drug) x Factor 2 (Training Stage)] | Training Stage: F(1, 9) = 31.04 *p*<.001 ; Drug: F(1, 9) = 0.8286, *p*=.3864, ns. ; Interaction: F(1, 3) = 4.788, *p*=.1165, ns. | None |
| 3E | Pre-PD: WT SAL v PSI | Average Session Time (min) | Two-way ANOVA  [Factor 1 (Drug) x Factor 2 (Training Stage)] | Training Stage: F(1, 9) = 15.63 *p*<.01 ; Drug: F(1, 9) = 3.651, *p*=.0884, ns. ; Interaction: F(1, 3) = 3.579, *p*=.1549, ns. | None |
| 3F | Pre-PD: WT SAL v PSI | % Correct (P+I+O) | Two-way ANOVA  [Factor 1 (Drug) x Factor 2 (Training Stage)] | Training Stage: F(1, 9) = 112.9, *p*<.0001 ; Drug: F(1, 9) = 3.596, *p*=.0904, ns. ; Interaction: F(1, 3) = 8.395, *p*=.0.0626, ns. | *P*D: SAL-*P*SI; *p* <.05 |
| 3G | Pre-PD: WT SAL v PSI | % Incorrect (P+I+O) | Two-way ANOVA  [Factor 1 (Drug) x Factor 2 (Training Stage)] | Training Stage: F(1, 9) = 67.82, *p*<.0001 ; Drug: F(1, 9) = 0.1172, *p*=.7400, ns. ; Interaction: F(1, 3) = 2.477, *p*=.2136, ns. | None |
| 3H | Pre-PD: WT SAL v PSI | % Omissions (P+I+O) | Two-way ANOVA  [Factor 1 (Drug) x Factor 2 (Training Stage)] | Training Stage: F(1, 9) = 3.905, *p*=.0795, ns. ; Drug: F(1, 9) = 2.999, *p*=.1174 ns. ; Interaction: F(1, 3) = 1.792, *p*=.2731, ns. | None |
| 3i | Pre-PD: WT SAL v PSI | % Correct (PD sessions only) | Two-way ANOVA  [Factor 1 (Drug) x Factor 2 (Session)] | Session: F(6, 54) = 10.95, *p*<.0001 ; Drug: F(1, 9) = 10.71, *p*<.01. ; Interaction: F(6, 34) = 0.5872, *p*=.7381, ns. | First session: SAL-*P*SI; *p*<.05 |
| 3j | Pre-PD: WT SAL v PSI | % Incorrect (PD sessions only) | Two-way ANOVA  [Factor 1 (Drug) x Factor 2 (Session)] | Session: F(6, 54) = 12.12, *p*<.0001 ; Drug: F(1, 9) = 6.719, *p*<.05. ; Interaction: F(6, 34) = 0.7951, *p*=.5803, ns. | None |
| 3L | Pre-R1: WT SAL v PSI | # Days to criterion | Two-way ANOVA  [Factor 1 (Drug) x Factor 2 (Training Stage)] | Training Stage: F(1, 7) = 7.896, *p*<.01 ; Drug: F(1, 7) = 0.09036, *p*=.7664, ns. ; Interaction: F(1, 2) = 0.09036, *p*=.7664, ns. | None |
| 3M | Pre-R1: WT SAL v PSI | # Sessions to criterion | Two-way ANOVA  [Factor 1 (Drug) x Factor 2 (Training Stage)] | Training Stage: F(1, 7) = 3.004, *p*=.1267, ns. ; Drug: F(1, 7) = 0.1829, *p*=.6817, ns. ; Interaction: F(1, 2) = 3.347, *p*=.2088, ns. | None |
| 3N | Pre-R1: WT SAL v PSI | # Trials to criterion | Two-way ANOVA  [Factor 1 (Drug) x Factor 2 (Training Stage)] | Training Stage: F(1, 7) = 6.563, *p*<.05 ; Drug: F(1, 7) =1.311, *p*=.2898, ns. ; Interaction: F(1, 2) = 3.503, *p*=.2021, ns. | None |
| 3O | Pre-R1: WT SAL v PSI | Average Session Time (min) | Two-way ANOVA  [Factor 1 (Drug) x Factor 2 (Training Stage)] | Training Stage: F(1, 7) = 7.257, *p*<.05 ; Drug: F(1, 7) =0.733, *p*=.4005, ns. ; Interaction: F(1, 2) = 5.628, *p*<.05. | None |
| 3P | Pre-R1: WT SAL v PSI | % Correct (P+I+O) | Two-way ANOVA  [Factor 1 (Drug) x Factor 2 (Training Stage)] | Training Stage: F(1, 7) = 8.351, *p*<.05 ; Drug: F(1, 7) =7.59, *p*<.05. ; Interaction: F(1, 2) = 4.125, *p*=.1793, ns. | R2: SAL-*P*SI; *p* <.05 |
| 3Q | Pre-R1: WT SAL v PSI | % Incorrect (P+I+O) | Two-way ANOVA  [Factor 1 (Drug) x Factor 2 (Training Stage)] | Training Stage: F(1, 7) = 10.62, *p*<.05. ; Drug: F(1, 7) =1.988, *p*=.2414, ns. ; Interaction: F(1, 2) = 2.582, *p*=.2493, ns. | None |
| 3R | Pre-R1: WT SAL v PSI | % Omissions (P+I+O) | Two-way ANOVA  [Factor 1 (Drug) x Factor 2 (Training Stage)] | Training Stage: F(1, 7) = 2.601, *p*=.1508, ns. ; Drug: F(1, 7) =2.151, *p*=.1859, ns. ; Interaction: F(1, 2) = 2.792, *p*=.2367, ns. | None |
| 3s | Pre-R1: WT SAL v PSI | % Correct (PD sessions only) | Two-way ANOVA  [Factor 1 (Drug) x Factor 2 (Session)] | Session: F(5, 35) = 8.535, *p*<.0001 ; Drug: F(1, 7) = 0.01705, *p*=.5534, ns. ; Interaction: F(5, 23) = 2.657, *p*=.8844, *p*<.05. | None |
| 3t | Pre-R1: WT SAL v PSI | % Incorrect (PD sessions only) | Two-way ANOVA  [Factor 1 (Drug) x Factor 2 (Session)] | Session: F(5, 35) = 11.22, *p*<.0001 ; Drug: F(1, 7) = 0.4524, *p*=., *p*=.4011, ns. ; Interaction: F(5, 23) = 0.1032, *p*=.9714, ns. | None |

### Supplementary Table 5. Statistical Summary of Experimental Outcomes for Figure 4

| **Figure #** | **Experimental Groups** | **Outcome Measure** | **Statistical Test** | **Main Statistics** | **Post-hoc Comparisons** |
| --- | --- | --- | --- | --- | --- |
| 4B | PD: Control vs ABA | # Days to criterion | Two-way ANOVA  [Factor 1 (Drug) x Factor 2 (ABA Exposure)] | ABA Exposure: F (1, 27) = 0.007390, p=.9321, ns.; Drug: F (1, 27) = 1.539, p=.2255, ns.; Interaction: F (1, 27) = 0.3332, p=.5686, ns. | None |
| 4C | PD: Control vs ABA | # Sessions to criterion | Two-way ANOVA  [Factor 1 (Drug) x Factor 2 (ABA Exposure)] | ABA Exposure: F (1, 27) = 0.04082, p=.8414, ns.; Drug: F (1, 27) = 0.3454, p=.5616, ns.; Interaction: F (1, 27) = 2.324, p=.1390, ns. | None |
| 4D | PD: Control vs ABA | # Trials to criterion | Two-way ANOVA  [Factor 1 (Drug) x Factor 2 (ABA Exposure)] | ABA Exposure: F (1, 27) = 0.2481, p=.6224, ns.; Drug: F (1, 27) = 0.2592, p=.6148, ns.; Interaction: F (1, 27) = 0.05541, p=..8157, ns. | None |
| 4E | PD: Control vs ABA | % Correct (P+I+O) | Two-way ANOVA  [Factor 1 (Drug) x Factor 2 (ABA Exposure)] | ABA Exposure: F (1, 35) = 4.482, p<.05; Drug: F (1, 35) = 0.9263, p=.3424, ns.; Interaction: F (1, 35) = 5.519, p<.05. | Control: SAL v PSI p<.05; PSI: Control vs ABA, p<.01 |
| 4F | PD: Control vs ABA | % Incorrect (P+I+O) | Two-way ANOVA  [Factor 1 (Drug) x Factor 2 (ABA Exposure)] | ABA Exposure: F (1, 35) = 3.032, p=.0904, ns; Drug: F (1, 35) = 0.7531, p=.3914, ns.; Interaction: F (1, 35) = 0.7479, p=.3930, ns. | None |
| 4G | PD: Control vs ABA | % Omissions (P+I+O) | Two-way ANOVA  [Factor 1 (Drug) x Factor 2 (ABA Exposure)] | ABA Exposure: F (1, 35) = 1.375, p=.2489, ns; Drug: F (1, 35) = 0.2491, p=.6208, ns.; Interaction: F (1, 35) = 3.158, p=.0843, ns. | None |
| 4H | PD: Control vs ABA | Average Session Time (min) | Two-way ANOVA  [Factor 1 (Drug) x Factor 2 (ABA Exposure)] | ABA Exposure: F (1, 29) = 0.04953, p=.8254, ns.; Drug: F (1, 29) = 1.431, p=.2413, ns.; Interaction: F (1, 29) = 1.484, p=.2330, ns. | None |
| *A2* | ABA: Pre-SAL vs Pre-PSI | Mean daily body weight loss | Independent T-Test | t(15) =0.5021, p=.6229, ns. | N/A |
| *A2* | ABA: Pre-SAL vs Pre-PSI | Mean daily food intake (g) | Independent T-Test | t(15) =0.03426, p=.9731, ns. | N/A |
| *A2* | ABA: Pre-SAL vs Pre-PSI | Mean daily total counts | Two-way ANOVA  [Factor 1 (Drug) x Factor 2 (ABA Stage)] | ABA Stage: F (1, 15) = 161.4, p<.0001; Pre-Treatment: F (1, 15) = 0.08435, p=.7755, ns.; Interaction: F (1, 15) = 1.126, p=.3053, ns. | None |
| 4J | R1: Control vs ABA | # Days to criterion | Two-way ANOVA  [Factor 1 (Drug) x Factor 2 (ABA Exposure)] | ABA Exposure: F (1, 27) = 3.058, p=.0917, ns.; Drug: F (1, 27) = 2.215, p=.1483, ns.; Interaction: F (1, 27) = 0.8652, p=.3605, ns. | Control: SAL v PSI, p=.0818; PSI: Control v ABA, p=.0621 |
| 4K | R1: Control vs ABA | # Sessions to criterion | Two-way ANOVA  [Factor 1 (Drug) x Factor 2 (ABA Exposure)] | ABA Exposure: F (1, 27) = 1.539, p=.2255, ns.; Drug: F (1, 27) = 3.434, p=.0748, ns.; Interaction: F (1, 27) = 2.586, p=.1195, ns. | None |
| 4L | R1: Control vs ABA | # Trials to criterion | Two-way ANOVA  [Factor 1 (Drug) x Factor 2 (ABA Exposure)] | ABA Exposure: F (1, 27) = 42.97, p<0.0001.; Drug: F (1, 27) = 0.14472, p=.7066, ns.; Interaction: F (1, 27) = 0.01374, p=.9075, ns. | SAL: Control vs ABA, p<.0001; PSI: Control vs ABA, p<.0001 |
| 4M | R1: Control vs ABA | % Correct (P+I+O) | Two-way ANOVA  [Factor 1 (Drug) x Factor 2 (ABA Exposure)] | ABA Exposure: F (1, 35) = 9.489, p<.01; Drug: F (1, 35) = 1.440, p=.2381, ns.; Interaction: F (1, 35) = 0.3565, p=.5543, ns. | SAL: Control vs ABA, p<.05; PSI: Control vs ABA, p=.0868 |
| 4N | R1: Control vs ABA | % Incorrect (P+I+O) | Two-way ANOVA  [Factor 1 (Drug) x Factor 2 (ABA Exposure)] | ABA Exposure: F (1, 35) = 3.442, p=.0720; Drug: F (1, 35) = 0.6050, p=.4419, ns.; Interaction: F (1, 35) = 0.08256, p=.7756, ns. | None |
| 4O | R1: Control vs ABA | % Omissions (P+I+O) | Two-way ANOVA  [Factor 1 (Drug) x Factor 2 (ABA Exposure)] | ABA Exposure: F (1, 35) = 9.858, p<.01; Drug: F (1, 35) = 1.459, p=.2352, ns.; Interaction: F (1, 35) = 0.06105, p=.8063, ns. | SAL: Control vs ABA, p<.05; PSI: Control vs ABA, p<.05 |
| 4P | R1: Control vs ABA | Average Session Time (min) | Two-way ANOVA  [Factor 1 (Drug) x Factor 2 (ABA Exposure)] | ABA Exposure: F (1, 24) = 12.58, p<.01.; Drug: F (1, 24) = 4.189, p=0.0518; Interaction: F (1, 24) = 0.2175, p=.6451, ns. | SAL: Control vs ABA, p<.05; PSI: Control vs ABA, p<.01 |

### Supplementary Table 6. Statistical Summary of Experimental Outcomes for Figure 5

| **Figure #** | **Experimental Groups** | **Outcome Measure** | **Statistical Test** | **Main Statistics** | **Post-hoc Comparisons** |
| --- | --- | --- | --- | --- | --- |
| 5C | R1: Control vs ABA | # Days to criterion | Two-way ANOVA  [Factor 1 (Drug) x Factor 2 (ABA Exposure)] | ABA Exposure: F (1, 26) = 0.2601, p=.6144, ns.; Drug: F (1, 26) = 0.05078, p=.8235, ns.; Interaction: F (1, 26) = 0.1025, p=.7514, ns. | None |
| 5D | R1: Control vs ABA | # Sessions to criterion | Two-way ANOVA  [Factor 1 (Drug) x Factor 2 (ABA Exposure)] | ABA Exposure: F (1, 26) = 0.3193, p=.5769, ns.; Drug: F (1, 26) = 0.3382, p=.5659, ns.; Interaction: F F (1, 26) = 2.392, p=.1340, ns. | None |
| 5E | R1: Control vs ABA | # Trials to criterion | Two-way ANOVA  [Factor 1 (Drug) x Factor 2 (ABA Exposure)] | ABA Exposure: F (1, 26) = 0.3000, p=.5886, ns.; Drug: F (1, 26) = 0.6189, p=.4386, ns.; Interaction: F (1, 26) = 1.589, p=.2187, ns. | None |
| 5F | R1: Control vs ABA | % Correct (P+I+O) | Two-way ANOVA  [Factor 1 (Drug) x Factor 2 (ABA Exposure)] | ABA Exposure: F (1, 26) = 2.586, p=.1199, ns.; Drug: F (1, 26) = 0.3618, p=.5527, ns.; Interaction:F (1, 26) = 0.03849, p=.8460, ns. | None |
| 5G | R1: Control vs ABA | % Incorrect (P+I+O) | Two-way ANOVA  [Factor 1 (Drug) x Factor 2 (ABA Exposure)] | ABA Exposure: F (1, 26) = 0.5867, p=.4506, ns.; Drug: F (1, 26) = 0.2114, p=.6495, ns.; Interaction:F (1, 26) = 1.397, p=.2478, ns. | None |
| 5H | R1: Control vs ABA | % Omissions (P+I+O) | Two-way ANOVA  [Factor 1 (Drug) x Factor 2 (ABA Exposure)] | ABA Exposure: F (1, 26) = 0.8122, ns.; Drug: F (1, 26) = 0.8440, , p=. ns.; Interaction:F (1, 26) = 1.219, p=.2478, ns. | None |
| 5I | R1: Control vs ABA | Average Session Time (min) | Two-way ANOVA  [Factor 1 (Drug) x Factor 2 (ABA Exposure)] | ABA Exposure: F (1, 26) = 4.938, p<.05; Drug: F (1, 26) = 0.2886, p=.5957, ns.; Interaction:F (1, 26) = 1.597, p=.2175, ns. | SAL: Control vs ABA, p<.05 |
| 5K | R2: Control vs ABA | % Correct (P+I+O) | Two-way ANOVA  [Factor 1 (Drug) x Factor 2 (ABA Exposure)] | ABA Exposure: F (1, 22) = 0.2062, p=.6542, ns.; Drug: F (1, 22) = 2.112, p=.1602, ns.; Interaction: F (1, 22) = 6.032, p<.05. | Control: SAL v PSI, p<.05 |
| 5L | R2: Control vs ABA | % Incorrect (P+I+O) | Two-way ANOVA  [Factor 1 (Drug) x Factor 2 (ABA Exposure)] | ABA Exposure: F (1, 22) = 0.2010, p=.6583, ns.; Drug: F (1, 22) = 0.02410, p=.8780, ns.; Interaction: F (1, 22) = 0.9563, p=.3388, ns. | None |
| 5M | R2: Control vs ABA | % Omissions (P+I+O) | Two-way ANOVA  [Factor 1 (Drug) x Factor 2 (ABA Exposure)] | ABA Exposure: F (1, 22) = 0.003273, p=.9549, ns.; Drug: F (1, 22) = 1.854, p=.1871, ns.; Interaction: F (1, 22) = 2.695, p=.1149, ns. | None |
| 5N | R2: Control vs ABA | Average Session Time (min) | Two-way ANOVA  [Factor 1 (Drug) x Factor 2 (ABA Exposure)] | ABA Exposure: F (1, 22) = 0.4737, p=.4985, ns; Drug: F (1, 22) = 0.2501, p=.6220, ns.; Interaction:F (1, 22) = 4.605, p<.05. | PSI: Control vs ABA, p<.05 |

### Supplementary Results

**Model comparisons**

The two generative learning models were compared to demonstrate that under BIC, the Rescorla–Wagner model provided the better fit in 86 of 87 sessions and in all 9 of 9 rats when session-wise BIC values were summed within animal. Mean model-comparison indices also favoured the RW model (mean rat-level BIC: RW = 1171.35, AI = 1306.42; mean rat-level AIC: RW = 1097.43, AI = 1183.22), indicating that the simpler RW account explained behaviour better than the active inference implementation.

**RW RL Model Results**

In the psilocybin group, the winning RW model showed descriptive shifts toward higher learning rate (Δα=+0.24, d=1.10, p=0.125), lower inverse temperature (Δβ=−0.50, d=−0.58, p=0.31), and lower initial value bias (ΔV0=−0.35, d=−1.16, p=0.125) from pre-drug to post-drug sessions**.** In saline controls, parameter changes were minimal (∣Δ∣≤0.03, ∣d∣≤0.11, p≥0.88).

**RL RW model parameter recovery**.

To assess whether the RW parameters were identifiable given the amount of data per rat, we performed a parameter-recovery analysis. For each fitted rat–session parameter set $\left( \alpha,\beta,V_{0} \right)$we simulated new choice data under the RW model, re-fit the same model, and correlated the true vs recovered parameters across all simulated data sets. Recovery was high for the learning-rate parameter (α: r ≈ 0.85) and moderate-to-high for the inverse temperature and bias parameters (β: r ≈ 0.66; V₀: r ≈ 0.61). Cross-parameter correlations were smaller in magnitude than the corresponding true–recovered correlations (e.g., modest trade-off between α and β), indicating that the model can reliably distinguish variation in the three parameters. These results suggest that the observed group differences in α and V₀ are unlikely to be artefacts of poor parameter identifiability.

#### Supplementary References

1. Anantharaman-Barr, H.G. and Decombaz, J., *The effect of wheel running and the estrous cycle on energy expenditure in female rats.* Physiol Behav, 1989. **46**(2): p. 259–63.

2. Verharen, J.P.H., Kentrop, J., Vanderschuren, L., and Adan, R.A.H., *Reinforcement learning across the rat estrous cycle.* Psychoneuroendocrinology, 2019. **100**: p. 27–31.

3. Cora, M.C., Kooistra, L. and Travlos, G., *Vaginal Cytology of the Laboratory Rat and Mouse: Review and Criteria for the Staging of the Estrous Cycle Using Stained Vaginal Smears.* Toxicol Pathol, 2015. **43**(6): p. 776–93.

4. Huang, K.*, et al.*, *Rapid, automated, and experimenter-free touchscreen testing reveals reciprocal interactions between cognitive flexibility and activity-based anorexia in female rats.* Elife, 2023. **12**.

5. Conn, K.*, et al.*, *Psilocybin restrains activity-based anorexia in female rats by enhancing cognitive flexibility: contributions from 5-HT1A and 5-HT2A receptor mechanisms.* Molecular Psychiatry, 2024. **10**: p.3291-3304.

6. Lopes, G. and Monteiro, P., *New Open-Source Tools: Using Bonsai for Behavioral Tracking and Closed-Loop Experiments.* Front Behav Neurosci, 2021. **15**: p. 647640.

7. Fisher, E.L.*, et al.*, *Psilocybin increases optimistic engagement over time: computational modelling of behaviour in rats.* Transl Psychiatry, 2024. **14**(1): p. 394.
